# The tRNA epitranscriptomic landscape and RNA modification enzymes in *Vibrio cholerae*

**DOI:** 10.1101/2025.10.15.682536

**Authors:** Léo Hardy, Virginie Marchand, Valérie Bourguignon, Quentin Thuillier, Cathy Dias, Evelyne Krin, Louna Fruchard, Dan Bar Yaacov, Didier Mazel, Yuri Motorin, Zeynep Baharoglu

## Abstract

Transfer RNAs (tRNAs) are central to protein synthesis, ensuring precise decoding of the genetic code by delivering aminoacids to the ribosome. Among all RNA species, tRNAs are the most heavily and diversely modified, with modifications playing critical roles in stability, folding, and function. Here, we present a comprehensive, isodecoder-level map of tRNA modifications in the human pathogen *Vibrio cholerae*. This map was generated by chemical-based sequencing methods, comparing wild-type and deletion strains. By assigning specific tRNA modifications to their cognate enzymes, we defined a comprehensive modification landscape in *Vibrio cholerae* and confirmed species-specific features, such as the presence of a functional TrmK enzyme, largely restricted to Gram-positive bacteria. Additionally, we detected a modification at U55 that occurs independently of TruB. To assess the biological significance of these modifications, we evaluated fitness under both standard conditions and subinhibitory antibiotic stress, and examined how modifications in the anticodon stem-loop region influence codon decoding efficiency and accuracy. Based on a comparative analysis of *E. coli* and *V. cholerae*, we discuss how species-specific differences in tRNA isodecoder gene repertoires may influence the functional impact and biological importance of tRNA modifications. This work provides the first experimentally validated, genome-wide map of tRNA modifications in *V. cholerae*, serving as a reference for future research into RNA modifications, translation regulation, and pathogen biology.

**Author summary:** This study charts the first genome-wide map of transfer RNA (tRNA) modifications in the cholera pathogen, *Vibrio cholerae*, revealing how chemical marks on tRNAs shape translation and stress responses. Using complementary chemical sequencing methods and a panel of targeted gene deletions, we assigned specific modifications to their enzymes across individual tRNA isodecoders. This integrative approach validates conserved features (e.g., Ψ55 and T54), and specific ones, such as an active TrmK that installs m¹A22 despite being considered largely restricted to Gram-positive bacteria, and uncovers enzyme interplay among dihydrouridine synthases. By testing mutant strains in standard and sub-inhibitory antibiotic conditions, we show that several modifications are dispensable for basal growth but become critical under proteotoxic stress, influencing fitness and translation accuracy, including stop-codon readthrough. Codon-specific reporter assays further demonstrate that modifications at wobble position 34 and at position 37 modulate decoding of distinct codon families, linking epitranscriptomic changes to gene expression programs. Comparative analysis with *Escherichia coli* suggests that species-specific tRNA isodecoder repertoires tune the functional impact of modifications. Our map provides an additional reference for studying RNA modification biology in pathogens and how it contributes to stress adaptation and virulence.

## Introduction

The understanding of the physiological roles of tRNAs has significantly advanced through the study of their extensive post-transcriptional modifications [1]. These modifications, introduced by highly specific tRNA-modifying enzymes, affect all four nucleotide types and encompass a diverse array of chemical changes, including base and sugar modifications such as methylation (e.g., m^6^A, m^5^C), pseudouridylation (Ψ), thiolation (s^4^U), and dihydrouridylation (D) [2]. In Gram-negative bacteria, tRNAs are typically modified at an average of eight positions per molecule, with modified nucleotides comprising approximately 10% of the total sequence [3]. These modifications are crucial for tRNA stability, folding, and decoding efficiency, highlighting their essential role in fine-tuning gene expression (for a recent review: [4]).

To date, over 100 distinct tRNA modifications have been identified across the three domains of life, including 18 universal modifications conserved in all domains [5, 6]. In the model Gram-negative bacterial organism *Escherichia coli*, 33 different modifications have been mapped across its 47 tRNA species. The number of dedicated tRNA-modifying enzymes is also remarkably high: for instance, among the ∼4,300 genes in the *E. coli* genome, 59 are involved in tRNA modification pathways, representing more than 1% of its coding capacity [7]. These modifications affect various aspects of tRNA biology depending on their position within the molecule: those located in the D-arm or T-arm primarily contribute to structural stability [8–10], while those in the anticodon stem loop modulate translational fidelity [11]. The anticodon stem loop is the most extensively modified region of the tRNA [12], particularly at position 34, the wobble position that pairs with the third base of the codon, and at position 37, which lies immediately 3′ to the anticodon.

While most tRNA modification genes can be inactivated without causing major phenotypic changes under optimal growth conditions, their absence can lead to a wide array of phenotypes, particularly under stress [12]. These include altered stress tolerance [13–21], changes in biofilm formation [22], modifications in metal transport [23], and impacts on virulence [24, 25] or motility [26, 27]. Moreover, some modifications influence the formation of others, a phenomenon known as modification circuits, suggesting a highly coordinated layer of epitranscriptomic regulation [28–30].

To uncover the functional consequences of tRNA modifications, it is essential to characterize their distribution across tRNA isodecoders and identify the enzymes responsible for their synthesis. The emergence of high-throughput sequencing-based technologies has now made possible to detect RNA modifications with single-nucleotide precision, even in organisms where traditional biochemical tools are limited [31–36]. The analysis of tRNA modifications has been performed in an increasing number of bacterial species, such as *Pseudomonas aeruginosa* [37], *Thermus thermophilus* [38], *Streptomyces albidoflavus [39]*, *Bartonella* spp. [40], *Staphylococcus aureus [41]*, *Methanocaldococcus jannaschii* [42], and thermophilic aerobic bacilli [43]. However, an exhaustive identification of the corresponding modifying enzymes, which is crucial for the study of associated phenotypes, has been performed only in a limited number of organisms, including *Escherichia coli* [7], *Mycoplasma capricolum [44]*, and *Bacillus subtilis [45]*.

In the Gram-negative pathogen *Vibrio cholerae*, our model organism and the causative agent of cholera, several unique tRNA modifications have been identified, including a cytosine to pseudouridine (C-to-Ψ) editing mechanism [46], and acetylation of acp^3^U [47], highlighting its potential for novel RNA processing pathways [48]. However, a comprehensive analysis of the full tRNA modification landscape and the systematic identification of the corresponding enzymes has never been undertaken. Recent studies from our laboratory have shown that tRNA modifications play a key role in *V. cholerae*’s adaptation to environmental stresses, including antibiotic exposure [20, 49].

In this study, we leveraged a combination of complementary deep-sequencing approaches, to comprehensively map the tRNA modification landscape of *V. cholerae*. These analyses were coupled with a systematic characterization of 22 individual gene deletion mutants targeting predicted or annotated tRNA-modifying enzymes.

Altogether, our integrated approach reveals the diversity and specificity of tRNA modifications in *V. cholerae*, and indicates stress-responsive modifications that may contribute to adaptive translation. These findings provide insights in bacterial tRNA modifications diversity and further establish *V. cholerae* as a relevant model for studying bacterial epitranscriptomics. They also underscore the need for a global characterization of tRNA modifications in additional organisms, in order to better understand their contribution to bacterial physiology, pathogenicity, and stress responses.

## Results

### Identification of annotated or putative genes of *V. cholerae* tRNA modification enzymes

Based on sequence homology and available annotations, we have first identified genes encoding known and potential tRNA modification enzymes in *V. cholerae*. **Table 1**.

**Table 1.**
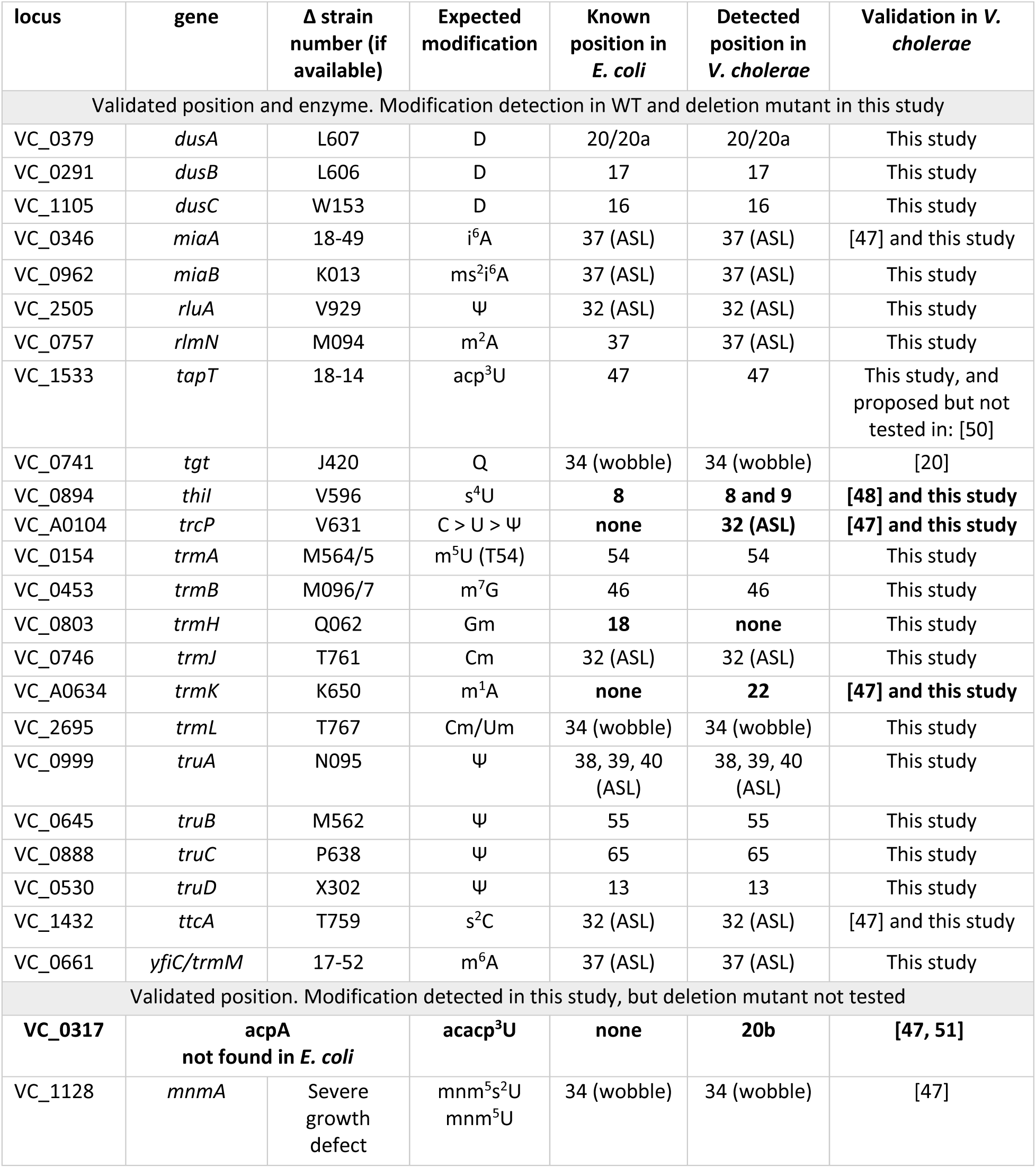
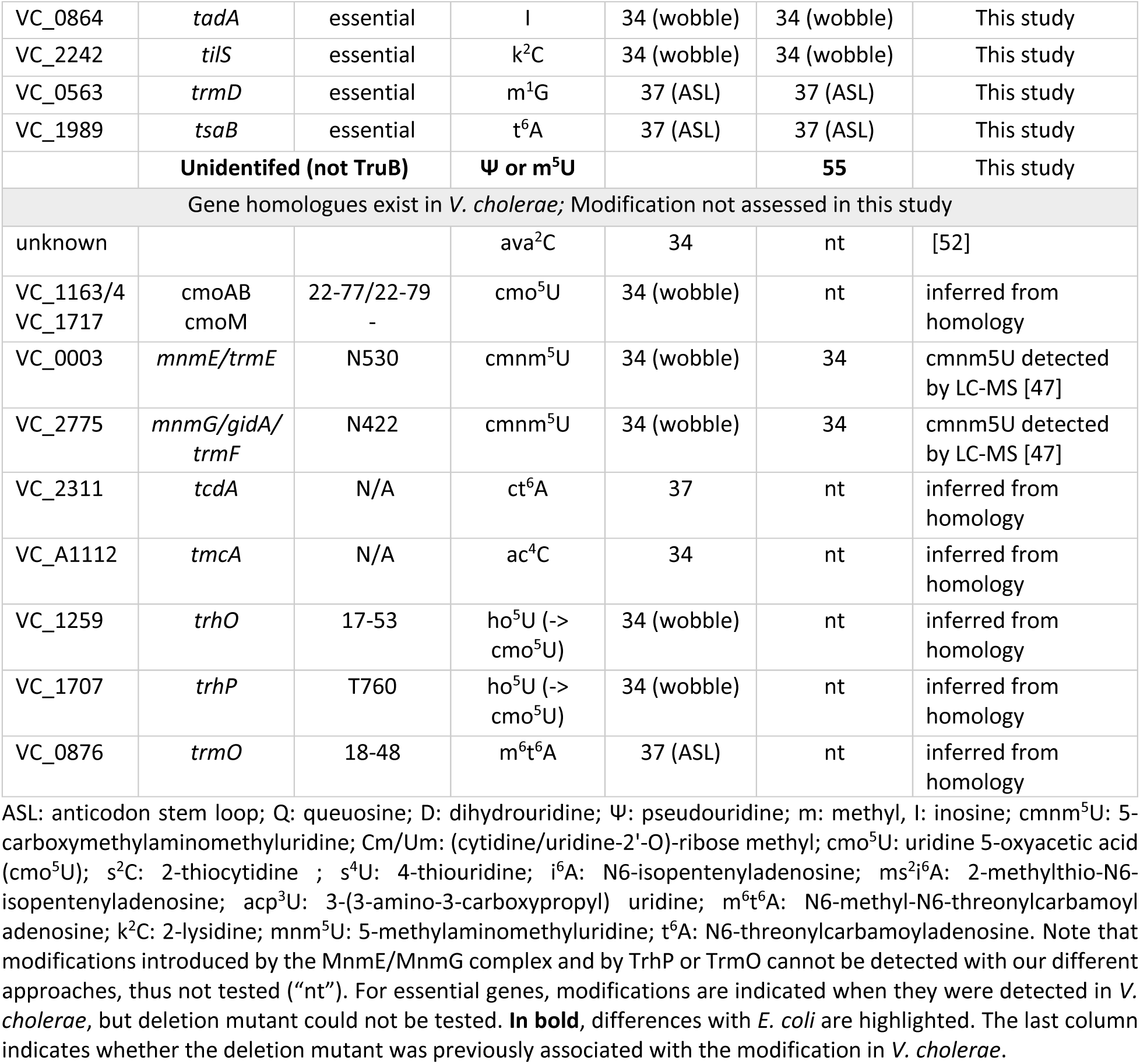
tRNA modification genes and associated modifications in *V. cholerae*

To comprehensively map tRNA modifications and identify the corresponding enzymes in *Vibrio cholerae*, we constructed and analyzed 23 individual tRNA modification gene deletion strains (**Table 1**). Several genes—*tadA*, *tsaB*, *tilS*, and *trmD*—are essential in *V. cholerae* (as determined by transposon insertion sequencing [19] and could not be deleted. Similarly, deletion of *mnmA* resulted in a severe growth defect, precluding further phenotypic analysis.

### Mapping of modified x^6^A with GLORI

In bacterial tRNAs, N^6^-modified adenosine residues (collectively referred to here as x^6^A) typically occur at position 37 within the anticodon loop . The GLORI protocol [53] enables reliable detection of those, including m^6^A, but also i^6^A (catalyzed by MiaA in *E. coli*), t6A (TsaB) and their derivatives, which are commonly found in bacterial tRNAs. GLORI detects x^^6^^A by selectively deaminating unmodified adenosines to inosines (read as G in sequencing) while modified adenosines resist conversion. Thus, unmodified adenosines appear as A→G substitutions, while modified adenosines are retained as A in the sequencing signal. While GLORI does not distinguish between different types of x^^6^^A modifications, known sequence contexts can aid interpretation: i^^6^^A and its derivatives usually occur in an AA*A context (with the middle A* being modified), whereas t^6^A is typically found in a UA*A motif. m^6^A and m^6^t^6^A (methylated by TrmO) are rare in tRNAs and have only been reported in a single tRNA species in *E. coli* [54]. Note that presence or absence of the m^6^ on m^6^t^6^A cannot be detected using GLORI as long as the essential t^6^A modification (formed by TsaB) is present.

GLORI analysis of *V. cholerae* tRNAs revealed multiple modification signals at position A37, which can be attributed to t^^6^^A_₃₇_ (or m^^6^^t^^6^^A₃₇), i^^6^^A_₃₇_ (or ms²i^^6^^A_₃₇_), and m^^6^^A (see **Figure 1**). First, we annotated i^^6^^A_₃₇_ signals that disappeared in isodecoders from the ΔVC_0346 (*miaA*) mutant (**Figure 1A**). We also detected a t^^6^^A signals **(Figure 1B**) that did not disappear in the Δ*miaA* strain and thus was attributed to TsaB, whose function cannot be tested by deletion because it is essential in *V. cholerae*. A single m^^6^^A modification was detected in tRNA^Val_CAA^ and depends on the presence of VC_0661, which shares 39 % sequence identity with *E. coli* YfiC (**Figure 1C**). We also detected a weak signals at position A37 in tRNAs Arg^ACG^, Asp^GTC^ and His^GTG^, which disappeared in the Δ*rlmN* (VC_0757) mutant (**Figure S1**). RlmN catalyzes the m²A37 modification, and the *E. coli* and *V. cholerae* RlmN proteins share 74 % sequence identity over 96 % of their length. It is plausible that m^2^A methylation also partially protects A from deamination in GLORI protocol, we therefore annotated these substoichiometric signals as m^²^A37 in these tRNAs. All modifications detected by GLORI are summarized in **Figure 1D**.

**Figure 1:**
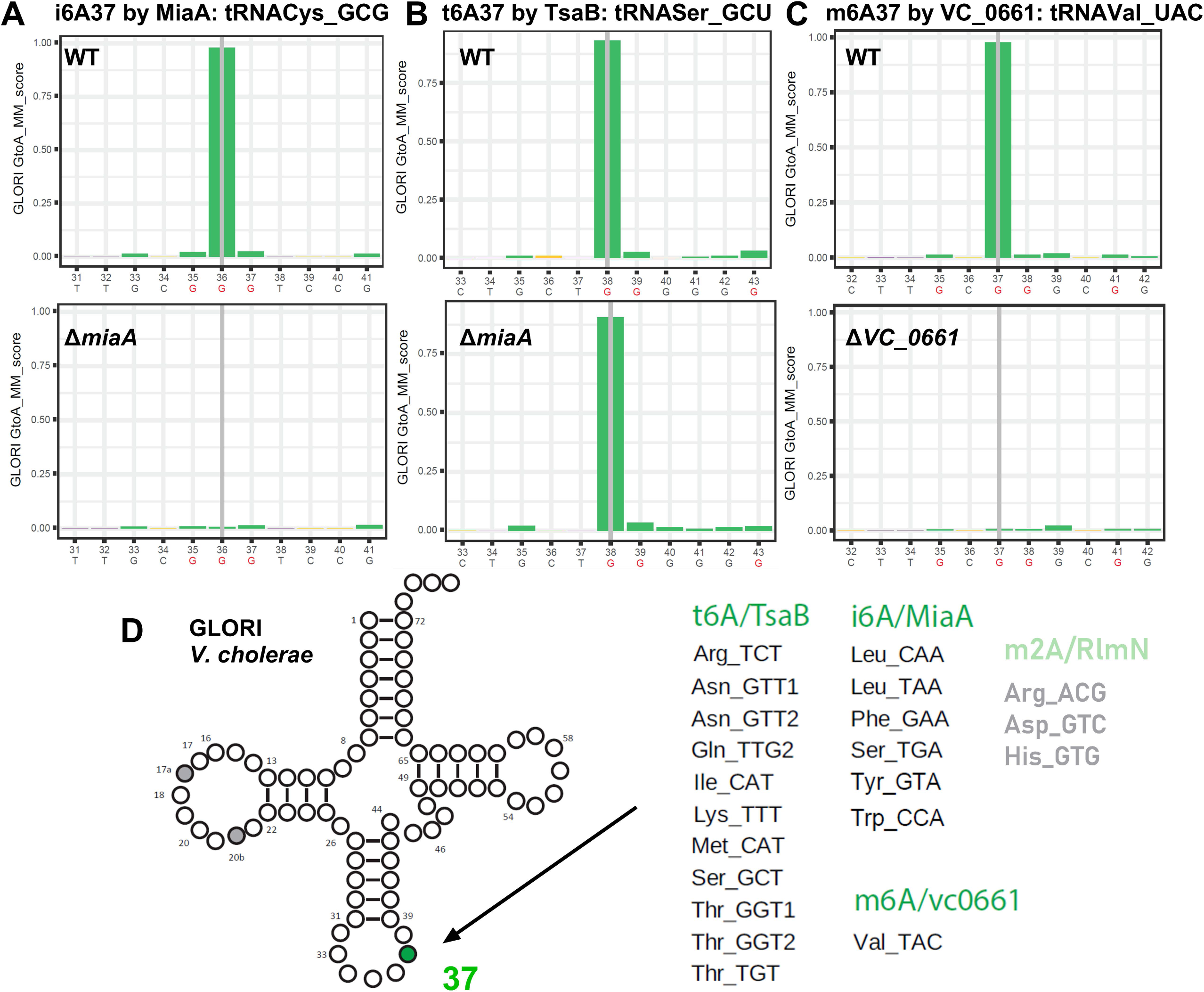
Mapping of modified x6A with GLORI. Histograms display the A-to-G deamination rate (y-axis) at the indicated nucleotide positions (x-axis). A→G substitutions indicate unmodified adenosines, whereas retention of A (high G-to-A score) denotes the presence of a modification. A: Detection of i^6^A37 made by MiaA in WT and absence of the signal in ΔmiaA. tRNA^Cys_GCG^ is shown as an example. B: Example for the detection of t6A made by TsaB in WT and ΔmiaA strains tRNA^Ser_GCT^. Note that tsaB is essential and the deletion mutant could not be tested. C: Detection of m6A in WT and the absence of the signal in the deletion mutant for VC_0661. tRNA^Val_UAC^ is shown as an example. G residues in tRNA resulting from A-to-I(G) conversion are shown in red. D: tRNA cloverleaf showing all modifications at position 37 detected by GLORI and the identity of isodecoders, including t^6^A, i^6^A and m^6^A. Note that the signal was low for the RlmN-dependent signal (in grey).

### Mapping of m^7^G/D/ho^5^C by AlkAnilineSeq

The AlkAnilineSeq method [33, 36] detects several RNA base modifications (including 7-methylguanosine (m^⁷^G), 3-methylcytosine (m^³^C), dihydrouridine (D), and the rare 5-hydroxycytidine (ho^^5^^C)) by selectively fragilizing and then aniline-cleaving the RNA at modified nucleotides, producing characteristic reverse-transcription stops or cDNA truncations for high-resolution mapping. Thus, cleavage occurs at modified nucleotides, generating a signal that is absent when the modification is missing.

Hydroxylated ho^5^C has so far been identified only in bacterial ribosomal RNA [55] and is not expected to be present in tRNA species. Likewise, m³C is not expected to occur in bacterial tRNAs. Consistent with this, AlkAnilineSeq analysis of *V. cholerae* tRNAs revealed no notable cytosine-derived signals.

In bacteria, 7-methylguanosine at position 46 (m^⁷^G46) is generally catalyzed by the TrmB methyltransferase. As predicted, many *V. cholerae* tRNAs contain m^⁷^G46 (**Figure 2A**). This modification was unambiguously attributed to VC_0453 TrmB, as the corresponding AlkAnilineSeq signal entirely disappeared in the *ΔtrmB* strain.

**Figure 2:**
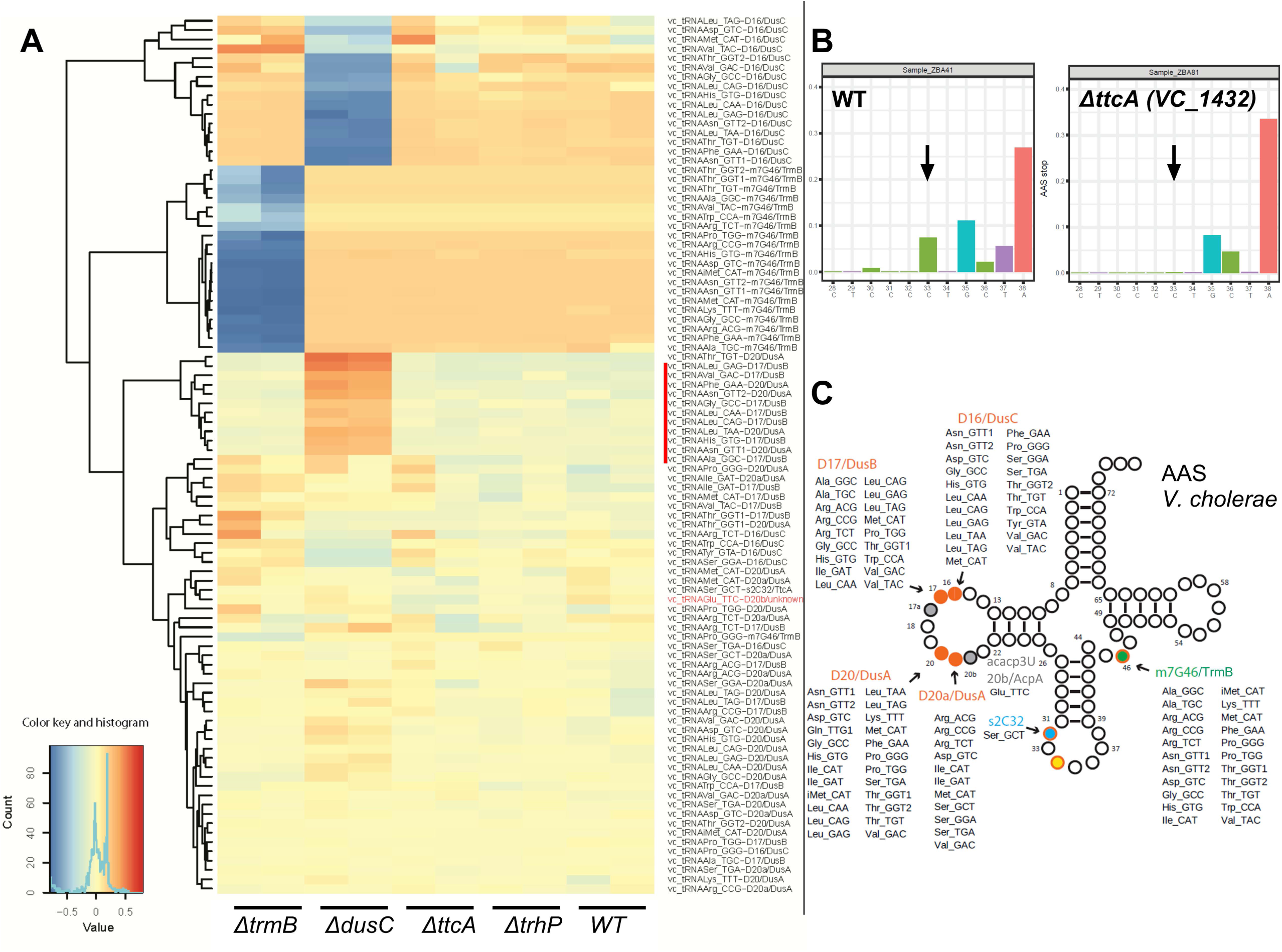
Mapping of m^7^G/D/ho^5^C by AlkAnilineSeq. A: Heatmap of detected modifications in WT, *ΔtrmB, ΔdusC, ΔttcA* and *ΔtrhP*. Differential heatmap, normalized to the average for all samples and the corresponding color key are shown. Blue indicates absence of modification while orange means increase. Isodecoders with increased D17 and D20 in the absence of *dusC* are indicated by a red vertical line. The tRNA^Glu_TTC^ U20b signal (acacp^3^U20) is indicated in red. **B:** Detection of s^2^C32 (arrow) in tRNA^Ser_GCU^ from WT and the absence of the signal in the deletion mutant for VC_1432. Aniline cleavage at modified nucleotide produces a detectable signal that is absent when the modification is lacking. **C:** tRNA cloverleaf showing all modifications detected by AlkAnilineSeq and the identity of isodecoders.

Another common tRNA feature is the presence of multiple dihydrouridine (D) residues in the D-loop, typically located between positions 16 and 20a. In bacteria, these modifications are introduced by dihydrouridine synthases (Dus), which belong to at least three distinct families: DusA, DusB, and DusC [56]. Functional annotation of these enzymes cannot be reliably done based on sequence homology alone, and deletion mutants are necessary for confident assignment of their activity and specificity [57]. Unlike the nearly stoichiometric nature of m^⁷^G, D modifications are typically substoichiometric and may be subject to regulation [58]. Using AlkAnilineSeq, we analyzed tRNAs from wild-type *V. cholerae* as well as ΔVC_0379 *dusA*, ΔVC_0291 *dusB*, and ΔVC_1105 *dusC* mutants. The results revealed a diverse set of tRNA substrates for each Dus enzyme, with D20 and D20a assigned to DusA, D17 to DusB, and D16 to DusC, like in *E. coli* (**Figure 2A**). Notably, deletion of *dusC* led to an increased signal at positions D17 (DusB target) and D20 (DusA target) in a subset of *V. cholerae* tRNAs (**Figure 2A**), indicating potential compensatory effects or cross-regulation among *V. cholerae* Dus enzymes.

Additionnaly, in tRNA^Glu_UUC^, a strong U-derived AlkAnilineSeq signal was observed at position U20b (**Figure 2A**), but this signal was unaffected by deletion of either *dusA*, *dusB* or *dusC*, or in the triple mutant, suggesting it arises from a distinct enzymatic activity. This signal is consistent with the previously reported acacp³U modification detected using LC-MS, catalyzed by AcpA and specific to *V. cholerae* [47, 51].

Finally, inspection of the AlkAnilineSeq profiles revealed a weak signal at position C32 in tRNA^Ser_GCU^. While this site is commonly modified to m³C in eukaryotic tRNAs [59, 60], it has been reported to carry a 2-thiocytidine (s²C) modification in bacteria [54, 61]. To test whether this weak signal originates from s^²^C, we analyzed tRNAs from a ΔVC_1432 (*ttcA*) strain, which lacks the enzyme predicted to catalyze s^²^C formation. Indeed, the AlkAnilineSeq signal at C32 was absent in the mutant, confirming the presence of s^²^C32 specifically in this *V. cholerae* tRNA (**Figure 2B**). Modifications detected using AlkAnilineSeq are summarized in **Figure 2C**.

### Mapping of 2’-O-methylations (Nm) residues by RiboMethSeq

RiboMethSeq [32, 34] maps 2′-O-methylated nucleotides by exploiting their resistance to alkaline cleavage: alkaline hydrolysis normally cleaves RNA randomly; however, the 3′ phosphodiester bond next to 2′-O-methylated (Nm) residues is relatively resistant to cleavage, as revealed at a protection at the N+1 nucleotide to Nm sites.

In *E. coli*, Nm at position 32 is introduced by TrmJ (typically as Cm or Um), whereas TrmL methylates position 34 (Nm). In *V. cholerae*, we identified genes encoding proteins homologous to *E. coli* TrmJ and TrmL: VC_0746 and VC_2695. RiboMethSeq analysis of *V. cholerae* WT tRNAs, combined with *ΔtrmJ* and *ΔtrmL* deletion mutants, allowed us to confidently identify a single 2’-O-methylated cytosine at position 32 (Cm32) in tRNA^Met_CAT^ (catalyzed by TrmJ), and two 2’-O-methylations at position 34 modifications: Cm34 in tRNA^Leu_CAA^ and Um34 in tRNA^Leu_TAA^ (both catalyzed by TrmL) (**Figure 3A**).

**Figure 3:**
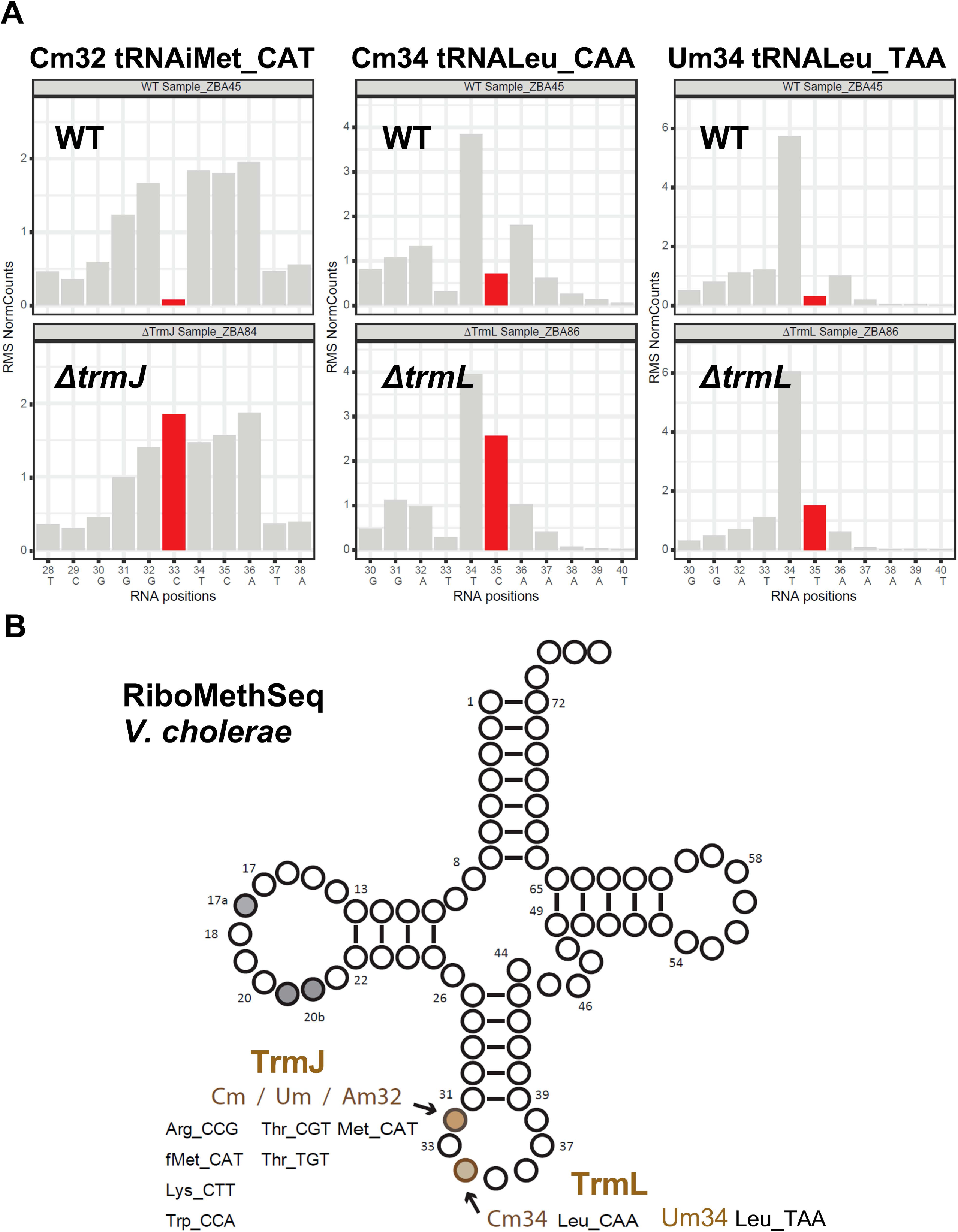
Mapping of 2’-O-methylations (Nm) residues by RiboMethSeq. A: Detection of Cm32, Cm34 and Um34 in WT and the absence of the protection signal in the deletion mutant for *trmJ* and *trmL*. Alkaline hydrolysis randomly fragments RNA, but the 3′-phosphodiester bond adjacent to Nm residues is more resistant to cleavage, producing a protection signal at positions containing Nm modifications (shown in red), this protection disappears in the mutant. Examples are shown. B: tRNA cloverleaf showing all modifications detected by RiboMethSeq and the identity of isodecoders.

In *E. coli*, TrmH catalyzes the 2′-O-methylation of guanosine at position 18 (Gm18) in the D-loop. In *V. cholerae*, however, no clear RiboMethSeq protection signal corresponding to Gm18 was detected, indicating that this site is unmodified, in contrast to *E. coli*. Furthermore, none of the protected residues detected in *V. cholerae* tRNAs exhibited altered cleavage patterns in the *ΔtrmH*-*like* (VC_0803) mutant strain, suggesting that this candidate enzyme may not act on tRNA and may possess different substrate specificity. **Figure 3B** summarizes modifications and substrates determined by RiboMethSeq.

### Mapping of pseudouridines (ψ), 5-methyl-uridine (m^5^U) and lysidine (k^2^C) by HydraPsiSeq and BID-Seq

Pseudouridine (ψ) residues in tRNAs were mapped using two orthogonal methods based on distinct chemistries. The first was HydraPsiSeq [35]. In HydraPsiSeq, pseudouridine (Ψ) bases are first chemically tagged by hydrazine so they can be distinguished from normal uridines. The RNA is then subjected to aniline cleavage and sequenced. Sites containing Ψ show a characteristic pattern, they resist cleavage and detected as protected U residues. HydraPsiSeq also detects 5-methyluridines (m^5^U) by their resistance to hydrazine cleavage and was also shown to detect lysidine (k^2^C), this conserved modified C at the tRNA position 34 is efficiently cleaved by hydrazine and thus can be distinguished from unmodified C. The second method, adapted from BID-Seq/PRAISE protocols [62, 63], where Ψ produces characteristic deletion signatures after bisulfite conversion under acidic conditions, enabling single-nucleotide resolution mapping of Ψ sites upon sequencing. The combination of HydraPsiSeq and BID-Seq allows for cross-validation of sites, significantly reducing both false positives and false negatives.

Four homologs of canonical tRNA-specific ψ-synthases were identified in the *V. cholerae* genome: VC_0999 for TruA (Ψ38/39/40 in *E. coli*), VC_0645 for TruB (Ψ55), VC_0888 for TruC (Ψ65), and VC_0530 for TruD (Ψ13). Our data confirm that these homologs in *V. cholerae* fulfill functions similar to those reported in *E. coli* (**Figure 4A**). TruA predominantly modifies position 39, with some tRNAs also carrying ψ38 or ψ40. TruC modifies two tRNAs at position 65, and TruD is responsible for a single Ψ13 site in tRNA^Glu_TTC^. Similar to *E. coli*, Ψ55 is detected in all tRNAs and is catalyzed almost exclusively by TruB. However, protection at U55 persisted in tRNA^Gln_TTG1^ and tRNA^iMet_CAT^ in *ΔtruB* (**Figure 4A**, red lines), indicating either the presence of an alternative enzyme or that the observed signal is not due to pseudouridine, but another hydrazine-resistant modification, like m^5^U, which is normally present at position 54 in bacterial tRNAs.

**Figure 4:**
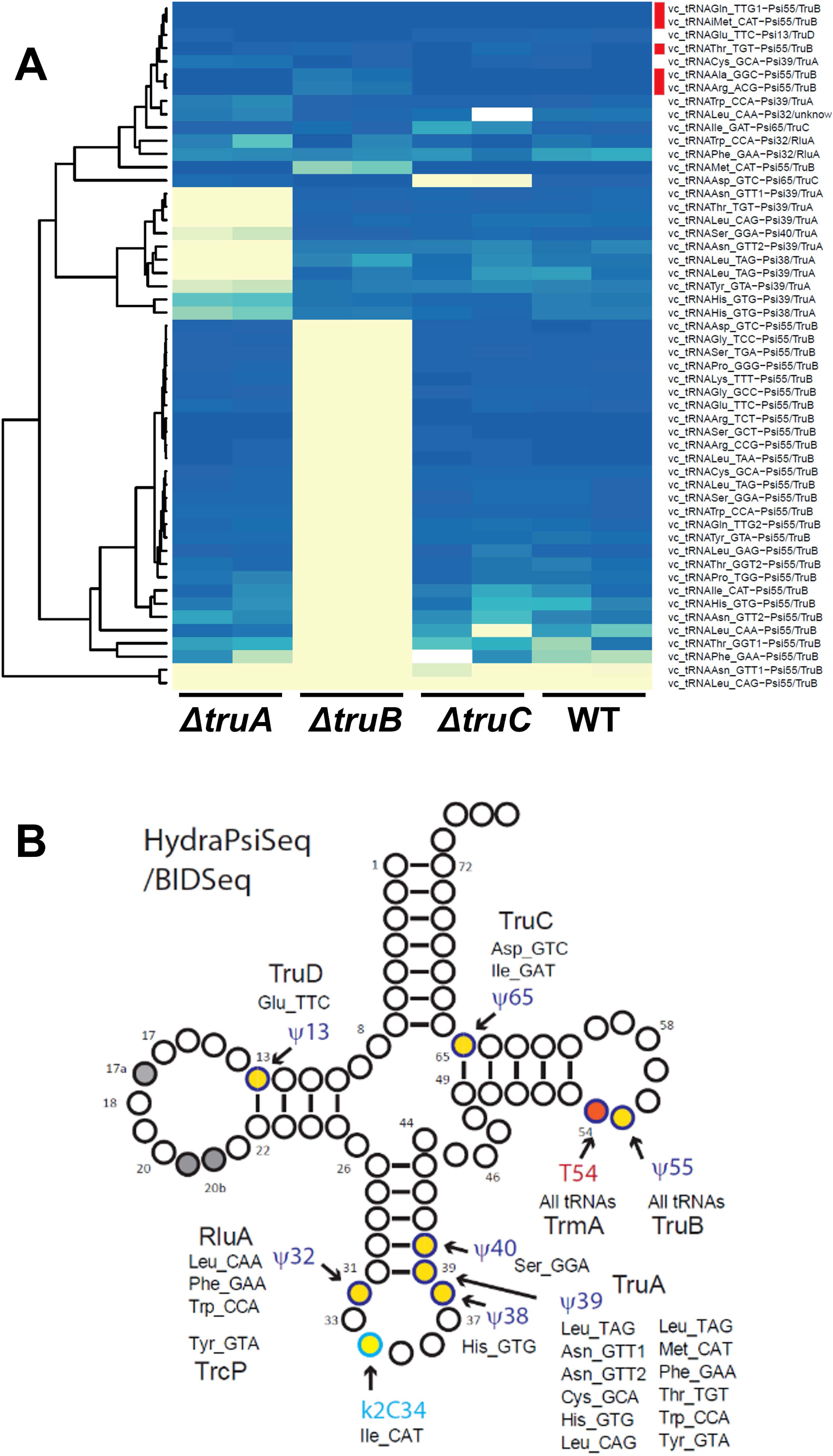
Mapping of pseudouridines (ψ) by HydraPsiSeq and BID-Seq. A. Heatmap of modifications detected by HydraPsiSeqin WT, *ΔtruA, ΔtruB, ΔtruC*, intensity of the blue colors indicates the strength of the HydraPsiSeq protection signal, which is related to the modification level. Yellow color corresponds to low signals from unprotected U residues. Isodecoders with invariable Ψ55 protection signal are indicated with a red vertical line.. B: tRNA cloverleaf showing all modifications detected by HydraPsiSeq and BID-seq and the identity of isodecoders.

RluA and RluF, dual-specificity pseudouridine synthases known to modify both tRNA and rRNA, and in the case of RluA, even mRNA [64] in *E. coli* [65], were also identified in *V. cholerae* as VC_2505 and VC_2223, respectively. Results show that *V. cholerae* RluA modifies tRNA^Trp^ and tRNA^Phe^ at position 32 (**Figure S2A**). Notably, although a RluF homolog is present in *V. cholerae*, no RluF-dependent ψ modifications were detected in any of the analyzed tRNAs.

Additionally, Ψ32 in *V. cholerae* tRNA^Tyr_GTA^ was previously shown to be generated through a unique two-step pathway, in which C-to-U RNA editing is followed by pseudouridylation catalyzed by TrcP (VC_A0104) [46]. We confirmed TrcP-driven formation of ψ32 in tRNA^Tyr^ and identified no additional tRNA substrates for this enzyme (**Figure S2B)**. Thus, ψ32 modification is installed through the combined action of RluA and TrcP in *V. cholerae*, depending on the tRNA.

Notably, HydraPsiSeq also detects 5-methyluridine (m^5^U), which is resistant to hydrazine cleavage. For the near-universal m^5^U54 (also called T54) modification in tRNA T-loop, *V. cholerae* possesses a TrmA-like enzyme: VC_0154 (**Table 1**). As expected, our data show that *V. cholerae* TrmA catalyze the formation of m^5^U54 across nearly all tRNA species (**FigureS2C**).

As anticipated from the results obtained previously for *E. coli* tRNA analysis [66], HydraPsiSeq analysis also revealed strong cleavage signal at position 34 of tRNA^Ile_CAT^, corresponding to lysidine, which is a cytidine whose C2 carbon is linked to an L-lysine moiety (k^²^C34) (**Figure S2D**) known to depend on TilS. The corresponding deletion mutant could not be tested because Tils (VC_2242; 45% sequence identity with *E. coli* TilS) is essential in *V. cholerae*.

Modifications detected using HydraPsiSeq and BIDseq are summarized in **Figure 4B. Mapping of m^5^C/m^4^C by RNA Bisulfite sequencing (BS-Seq)** Detection of cytosine base methylations, primarily 5-methylcytosine (m^^5^^C) and the much rarer 4-methylcytosine (m⁴C), is traditionally carried out using RNA bisulfite sequencing (BS-Seq), a method adapted from the well-established approach used for detecting m^5^C in DNA [67–69]. Unmodified cytosines are deaminated to uracils (read as T in DNA) upon bisulfite treatment under neutral or slightly alkaline pH, while m^^5^^C and m^⁴^C are resistant to this conversion and thus read as C in sequencing.

When we applied this technique to all *V. cholerae* tRNAs, we observed no significantly deamination-resistant cytosine residues, indicating the absence of detectable m^5^C or m⁴C modifications under the tested conditions. This observation is consistent with long-standing affirmation on the absence of m^5^C residues in bacterial tRNAs [54]. However, a recent study demonstrated the stress-induced appearance of m^5^C in *E. coli* tRNA^Tyr^, catalyzed by the RsmF ribosomal methyltransferase [70]. To investigate whether a similar stress-induced m^5^C formation occurs in *V. cholerae*, we re-analyzed tRNA extracted from cells exposed to 1 mM H₂O₂ for 10 minutes. tRNA BS-Seq analysis of these samples revealed no evidence of m^5^C modifications, neither in tRNA^Tyr^ nor in any other *V. cholerae* tRNA (**Figure S3A**). Thus, under the tested oxidative stress conditions, m^5^C formation does not appear to occur in *V. cholerae* tRNAs.

### Analysis of RT-signatures for mapping of RT-mismatching and RT-arresting tRNA modifications: s^4^U, acp³U, m^¹^G, ms²i^^6^^A, m^¹^A

Certain modified nucleotides induce base misincorporations (RT mismatches), nucleotide deletions or premature termination of cDNA synthesis (RT arrest), leaving reproducible RT signatures in sequencing data. RT-signature analysis uses these signatures to map the positions of tRNA modifications. However, such methods relying on natural reverse transcription (RT) signatures, which include RT stops, misincorporations, and deletions (“jumps”) at modification sites, are inherently less robust. First, these signals should be distinguished from single nucleotide variant, frequently present due to tRNA isoacceptors with closely related sequences. These single nucleotide variants may be variable and non-stoichiometric, reflecting differences in tRNA expression levels. Secondly, these RT-based signals are highly dependent on the properties of the reverse transcriptase used, and the only reliable control for confirming modification-specific effects is the use of deletion mutants lacking the corresponding modifying enzyme. As such, RT-signature analysis was used primarily as a complementary approach, and most data on deletion/misincorporation patterns were extracted from existing RiboMethSeq, AlkAnilineSeq, and HydraPsiSeq datasets.

Several common tRNA-modified residues are known to generate either simple reverse transcription (RT) signatures, such as mismatches, or more complex signatures involving combinations of mismatches, arrests, and jump events. First, 4-thiouridine (s⁴U), which is classically introduced by ThiI in bacterial tRNAs at position 8. Although s⁴U is known to base pair with adenine, it can also pair with guanine, resulting in a strong U→C transition in sequencing data. This signature can be detected in various datasets, but it is particularly prominent in bisulfite sequencing (BS-Seq) analysis. Inspection of *V. cholerae* tRNA bisulfite sequencing data (see above) revealed the presence of s⁴U8 in 22 *V. cholerae* tRNAs. Interestingly, a similar signal is also observed at position 9 in *V. cholerae* tRNA^Tyr_GTA^, consistent with previous report [48]. As expected, these signals were attributed to ThiI (VC_0894) (**Figure S3B).**

The modified nucleotide acp³U47 (uridine carrying a 3-amino-3-carboxypropyl side chain at its N3 position) is commonly found in bacterial tRNAs and is catalyzed by TapT. Because this chemical modification tends to block or slow reverse transcription (RT), the modification can in principle be detected by looking for mismatches in sequencing data. We found that this RT-based signal disappears after treating tRNAs with periodate (IO₄⁻) [20], indicating that acp³U is fully oxidized under these conditions, leaving only unmodified U47 molecules to be seen during sequencing. In addition, deleting the *tapT* gene (VC_1533) in *V. cholerae* also removed the RT-mismatch signal at U47, without periodate treatment. Together, these results confirm that TapT produces acp^³^U at position 47 and allowed us to map this modification in at least seven *V. cholerae* tRNAs (**Figure S3B**).

For selected deletion strains, we also performed RT-stop analysis to confirm the presence of specific tRNA modifications that block reverse transcription. In several tRNAs, tRNA^Pro_TGG^, tRNA^Leu_CAG^, and tRNA^Leu_GAG^, we detected characteristic reverse transcription stops and base-misincorporation signals at m¹G37, a modified guanosine made by the essential enzyme TrmD (encoded by the *V. cholerae* gene VC_0563). Likewise, strong RT arrest was seen at position 37 in tRNA^Leu_TAA^, tRNA^Trp_CCA^, and tRNA^Tyr_GTA^, where the hypermodified base ms²i^6^A37 is normally found (**Figure S3B**). This RT signature disappeared when the MiaB enzyme (VC_0962) was missing, consistent with the idea that the extra methylsulfur group in ms²i^6^A makes reverse transcription stall more strongly.

A TrmK-like enzyme (VC_A0634) in *Vibrio cholerae* was found to catalyze the formation of m¹A22 [71] **[47]**. RT-stop analysis showed a modification signal at position 22 in the D-loop of tRNA^Pro_TGG^ and tRNA^Tyr_GTA^; and this signal disappeared in the ΔVC_A0634 mutant. However, even in the wild-type strain the signal was weak and variable. Using the GLORI method (described above), we also detected a low, substoichiometric signal at the same A22 position in these tRNAs. As this site is known to carry the m^¹^A modification, the GLORI-positive A22 residues in *V. cholerae* tRNAs were identified as m¹A22, a modification previously confirmed in *V. cholerae* tRNAs by LC–MS analysis [47]. Our study confirms this role for TrmK and additionally reports the absence of TrmK-dependent modifications in other isodecoders. This finding supports that TrmK specifically modifies tyrosine and proline tRNAs. Interestingly, the methyltransferase TrmK, which catalyzes the formation of m¹A22, is typically found in Gram-positive Firmicutes (**Figure S4**). *V. cholerae* appears to be an unusual exception among Gammaproteobacteria, sharing this feature only with *Shewanella oneidensis*, which also encodes a TrmK-like gene [71].

### RNA-seq analysis for mapping A>I tRNA editing

Inosine (I), which commonly replaces A34 in ANN anticodon sequences, base pairs with cytosine and is thus read as guanosine (G) in sequencing data. In *E. coli* and many other organisms, the enzyme TadA catalyzes the deamination of adenosine to inosine at position 34 of tRNA^Arg_ACG^ [72]. In addition, tRNA^Leu_AAG^ was also recently shown to be edited by TadA in *Streptococcus pyogenes* [73]. We used our previously published RNAseq data [20] to assess A-to-I editing on tRNA^Arg^. RNA-seq analysis [74] confirmed the presence of inosine at position 34 in tRNA^Arg_ACG^, confirming that TadA function is conserved in this organism. The corresponding deletion mutant for *tadA* (VC_0864) cannot be tested as the gene is essential in *V. cholerae*. Interestingly, the frequency of inosine modification was found to increase under sub-lethal tobramycin treatment (**Figure S5A**), suggesting a possible role for this modification in stress adaptation or translational regulation during antibiotic exposure.

A comprehensive summary of all detected tRNA modifications and their associated enzymes is provided in **Figure 5**, with detailed isodecoder-specific data in **Figure 6**.

**Figure 5:**
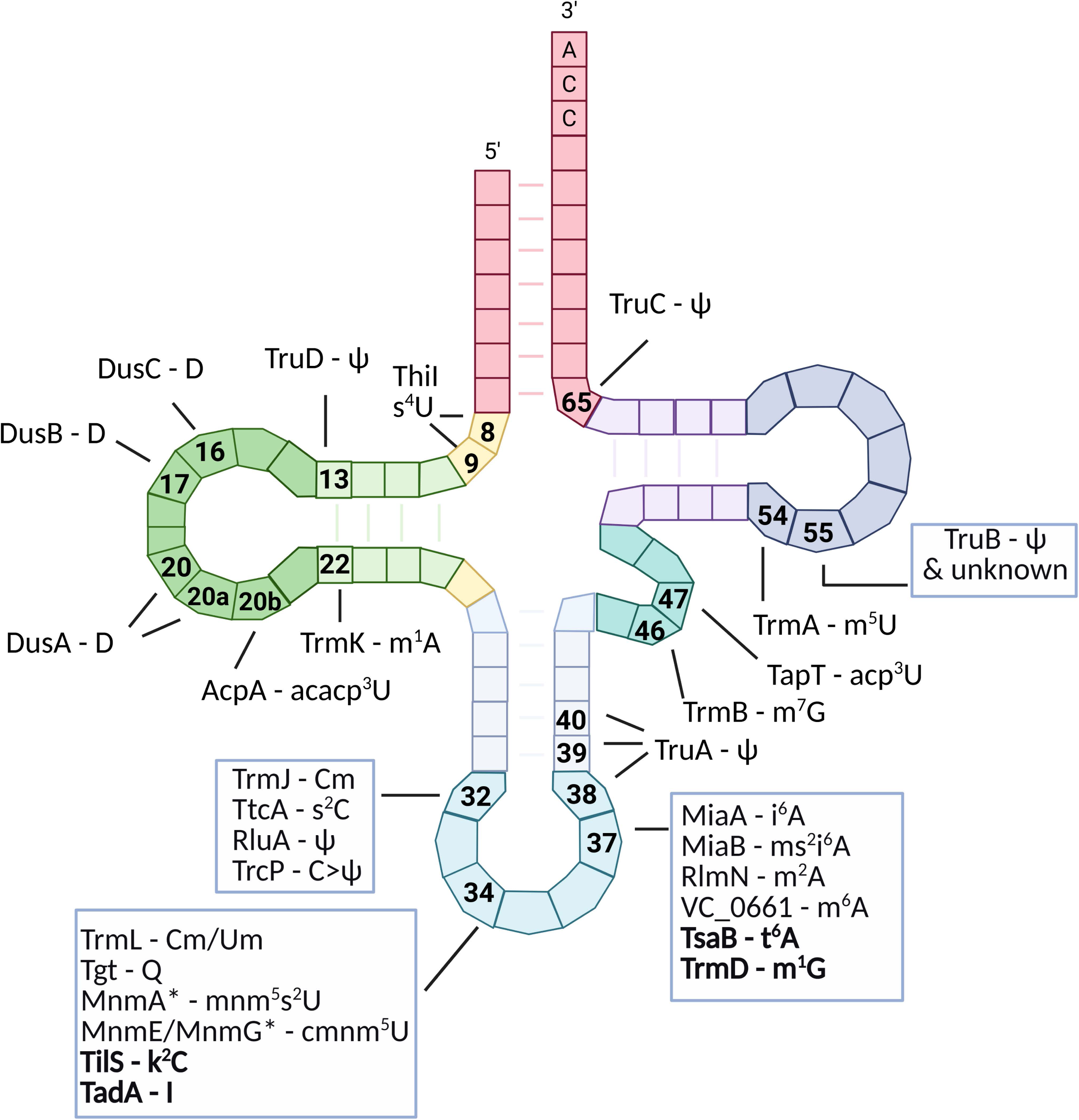
*V. cholerae* tRNA modifications overview. Secondary structure of a canonical bacterial tRNA annotated with RNA modifications detected in *Vibrio cholerae*. The tRNA is shown in its cloverleaf conformation. Post-transcriptional modifications identified in *V. cholerae* are mapped to their corresponding positions using standard abbreviations as in Table 1. The anticodon is centered at positions 34–36, position 34 indicated is the wobble base. Enzyme names are indicated next to their associated modifications; enzymes in bold are essential for *V. cholerae* growth and are presumed to catalyze the corresponding detected modifications. Modifications marked with an asterisk (*) are not detectable with current sequencing-based approaches but were detected previously by LC-MS and enzyme names are inferred from known modification pathways and presence of the homologous enzyme in *V. cholerae*.

**Figure 6:**
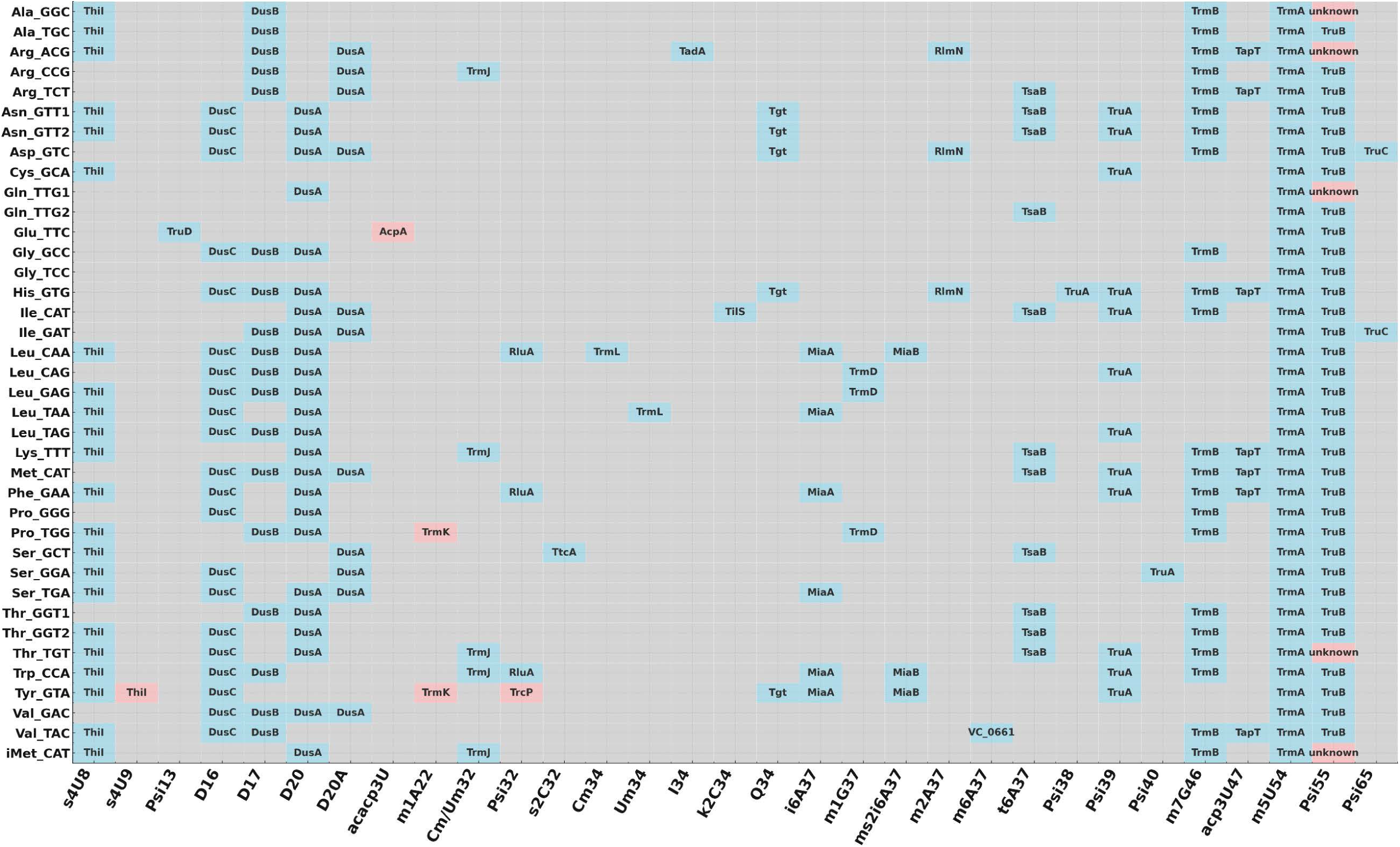
Overview of modifications of *V. cholera*e for all isodecoders. Each row represents a unique tRNA isoacceptor, labeled by its corresponding amino acid and anticodon. Columns denote individual modification sites ordered by nucleotide position. We indicate the presence of a modification in *V. cholerae*, with the name of the associated enzyme (or unknown) displayed when available. Grey cells indicate absence of modification. Pink cells indicate differences with *E. coli*.

### Impact of tRNA modification enzyme deletions in *V. cholerae* on fitness

Although several tRNA modification genes are essential, the majority of those that are nonessential show no detectable growth phenotype under standard laboratory conditions when deleted. In a previous study, we tested 13 *V. cholerae* deletion mutants for tRNA modification genes (*tgt, mnmG, dusA, dusB, miaB, truA, truB, truC, trmA, trmB, mnmE,* and *trmK*) under normal growth conditions and during proteotoxic stress induced by the aminoglycoside antibiotic tobramycin, revealing stress-specific phenotypes [19].

Here, we extended this analysis to newly constructed deletion strains, for position 34: Δ*trmL*, Δ*trhP*; for position 37: Δ*miaA*, Δ*trmO;* others in the anticodon loop: Δ*ttcA*, Δ*trcP*, Δ*trmJ* and the dual-function enzyme Δ*rluA; and* elsewhere: Δ*dusC*, Δ*truD*, Δ*tapT* (**figure S5B**).

Among these, only Δ*ttcA*, Δ*miaA*, and Δ*tapT* exhibited a growth defect under non-stress conditions, consistent with the general observation that tRNA modification deletions often do not impact basal fitness. However, several new mutants displayed distinct stress-specific phenotypes. Notably, deletions of *trhP*, *dusC*, and *rluA* conferred a growth advantage in the presence of tobramycin, suggesting potential roles in modulating stress response pathways. In contrast, Δ*ttcA*, Δ*tapT*, and Δ*miaA* showed a pronounced fitness defect under tobramycin stress, indicating that these modifications are particularly critical for survival under proteotoxic conditions. Further work is needed to dissect the underlying molecular mechanisms.

### Impact of tRNA modification enzyme deletions in *V. cholerae* on stop codon readthrough

We also quantified stop codon readthrough using previously described dual-reporter constructs [20, 75]. Our goal was to evaluate whether modifications within the anticodon loop have a stronger impact on translation accuracy at stop codons compared to modifications located elsewhere in the tRNA body (e.g., D-loop, T-loop, and other regions).

Our results indicate that the position of a modification, whether in the anticodon loop or outside of it, is not the primary determinant of its effect on stop codon readthrough. Instead, we observed that the UGA stop codon is the most strongly affected by the presence or absence of specific modifications, particularly under tobramycin (TOB) treatment. Within the anticodon loop, deletion of *tgt*, *miaB*, or *trmJ* led to a marked increase in stop codon readthrough in the presence of TOB, compared to the wild-type strain. In contrast, deletions of *trmL*, *trhP*, *ttcA*, *rluA* and *truA* reduced readthrough, particularly at UGA (**Figure S6A**).

For modifications outside the anticodon loop, increased readthrough was observed in the *ΔtruD* and Δ*trmB* mutants, while Δ*dusA*, Δ*dusB,* Δ*dusC*, and Δ*thiI* mutants showed decreased readthrough (**Figure S6A**). For *ΔtruC,* we observe increased readthrough at UAG but decreased at UGA. These findings demonstrate that a variety of tRNA modifications modulate translation accuracy in distinct ways, and highlight the need to investigate each modification individually to uncover its specific mechanism of action. We asked which tRNA-modification enzymes produce shared effects on stop-codon readthrough. We identified deletion mutants whose loss consistently increased or decreased readthrough across the three stop codons (UAA, UAG, UGA), presented as a Venn diagram (**Figure S6B**). We also examined which tRNA isodecoders are the most frequently shared substrates among enzymes that modulate readthrough. tRNA^Tyr_GTA^ appears most often, followed by tRNA^Trp_CCA^, with several tRNA^Leu/^tRNA^Pro/^tRNA^Phe^ isodecoders also recurrent. These patterns suggest that modifications on these tRNAs may contribute to translation accuracy; targeted functional studies will be needed to determine causal roles for specific modifications and sites.

### Codon-specific effects of anticodon loop modifications at position 34: impact of U34 modifications on G-ending codon decoding

tRNA modifications at the wobble base (position 34) and the adjacent purine (position 37) are the ones most strongly expected to influence codon recognition and translation accuracy. The nucleotide at position 34 pairs directly with the third base of the codon; chemical modifications there, introduced by enzymes such as Tgt, MnmEG, TrhP, TrhO, MnmA, TadA, TilS, and TrmL, fine-tune wobble pairing and help maintain the reading frame [76, 77]. Our analyses confirmed modification by Tgt and TrmL, and we detected modifications installed by the (nearly) essential TadA, TilS, and MnmA in *Vibrio cholerae*. Modifications mediated by MnmEG and TrhPO could not be tested with our methods. However, *V. cholerae* encodes the canonical MnmE/MnmG enzymes that install 5-carboxymethylaminomethyluridine (cmnm^5^U) at wobble U34, a reaction well established in bacteria and supported in *V. cholerae* by curated gene annotations and tRNA-modification profiling by mass spectrometry [47] (Table 1). ho^5^U has not yet been detected in *V. cholerae*, but its biosynthetic pathway remains plausible given the presence of TrhPO homologues [78].

To assess the codon-specific impact of tRNA modifications at positions 34 and 37, we employed our fluorescent reporter system, previously validated in Δ*tgt* (queuosine at position 34) [20]. Briefly, we inserted a stretch of three identical codons in tandem at the 5′ end of the GFP coding sequence (e.g., ATG_(start)_-TAC-TAC-TAC for tyrosine), so that the relative fluorescence of each reporter in a Δ*mutant* versus wild type background provides a readout of codon-specific decoding efficiency. For this analysis, we selected mutants lacking *trhP* and *mnmE*, as these genes are non-essential, and show no visible growth defect under standard conditions.

To probe the impact on decoding of U34 modifications, we focused on codons that vary at the third (wobble) position, pairing with U34 in the tRNA anticodon. Since both TrhP and MnmE modify U34, we compared decoding efficiency of A-ending versus G-ending codons across several codon families: Ser (TCA/TCG), Leu (TTA/TTG, CTA/CTG), Val (GTA/GTG), Pro (CCA/CCG), Gln (CAA/CAG), Arg (CGA/CGG, AGA/AGG), Thr (ACA/ACG), and Lys (AAA/AAG).

Our data reveal that deletion of either *trhP* or *mnmE* significantly impairs decoding of G-ending arginine codons (AGG and CGG), while A-ending counterparts (AGA and CGA) are largely unaffected (**Figure 7AB**).

**Figure 7.**
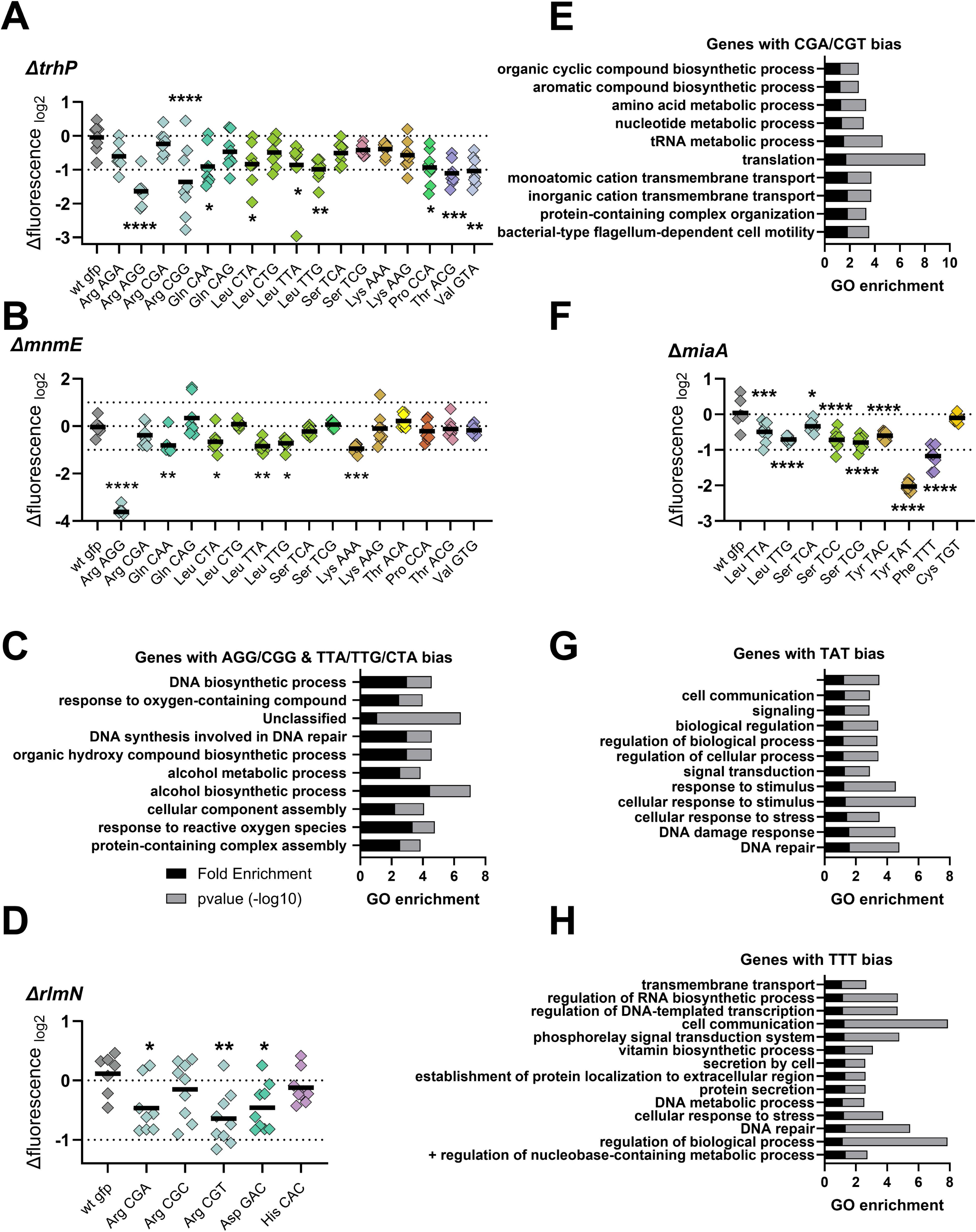
Codon-Specific Effects of selected modifications at positions 34 and 37. A and B: Codon-specific translation efficiency in WT and *Δmutant* strain using 3xcodon stretches inserted in GFP, as previously described (Fruchard et al, 2025). Y-axis represents the relative fluorescence of a given GFP in *Δmutant* over the same construct in WT. Each specified codon is repeated 3× within the coding sequence of the GFP (e.g. TACTACTAC). For multiple comparisons, we used one-way ANOVA. **** means p<0.0001, *** means p<0.001, ** means p<0.01, * means p<0.05. ns: non-significant. Number of biological replicates for each experiment: n=8, each dot represents one replicate. “wt gfp” is the native gfp (gfpmut3) without any stretch. **C:** Gene ontology enrichment analysis in *V. cholerae* of overrepresented gene categories with a usage bias for AGG,CGG, TTA, TTG, CTA. The bioinformatic workflow used for whole-genome codon bias calculations follows the approach described in [20] and detailed in Figure 4C of that study. **D:** Codon-specific translation efficiency in WT and *ΔrlmN*. **E:** Gene ontology enrichment analysis in *V. cholerae* of overrepresented gene categories with a usage bias for CGA/CGT. **F:** Codon-specific translation efficiency in WT and *ΔmiaA*. **G and H**: Gene ontology enrichment analysis in *V. cholerae* of overrepresented gene categories with a usage bias for TAT or TTT.

This supports the model wherein U34 modifications facilitate U-G wobble pairing without disrupting canonical U-A interactions. These findings align with previous work in *E. coli*, which demonstrated that MnmE-mediated modification stabilizes U–G pairing at NNG codons [79], and TrhPO’s impact on decoding of GCG and UCG [78]. Additionally, both mutants exhibit reduced decoding efficiency of leucine codons TTA, TTG, and CTA.

To explore the functional significance of these codon biases, we leveraged our existing codon usage datasets for *V. cholerae* [20] (and doi:10.5281/zenodo.6875293). We pooled genes enriched for the affected codons (Arg AGG/CGG and Leu TTA/TTG/CTA), generating a list of 854 genes out of 3781 annotated in the *V. cholerae* genome. Gene ontology (GO) enrichment analysis using the PANTHER database revealed overrepresentation of pathways related to protein complex assembly, response to reactive oxygen species, alcohol metabolism, and DNA repair (**Figure 7C**). This suggests that TrhP-and MnmE-dependent modifications may play a regulatory role in stress response through codon-biased translation.

Notably, 18 transcription factors were found to preferentially use these codons, including *oxyR* (VC_0732), a key regulator of the oxidative stress response (**Figure S7**). These findings support a model in which codon usage, coupled with dynamic tRNA modification, modulates stress-responsive gene expression in *V. cholerae*.

### Codon-specific effects of anticodon loop modifications at position 37

Position 37, immediately 3′ of the anticodon, often carries modifications such as m^1^G37 (TrmD), i^6^A37 (MiaA), m^2^A (RlmN), or t^6^A37 (TsaB), which stabilize codon-anticodon interactions and prevent frameshifting [80, 81]. We confirmed these modifications in WT and deletion mutant except for TrmD and TsaB which are essential. To examine the functional consequences of lacking A37 modifications, we evaluated decoding efficiency in the mutant lacking *miaA* or *rlmN*, using codons recognized by the corresponding tRNA isodecoders. Note that *ΔrlmN* shows no growth defect while Δ*miaA* exhibits a mild fitness reduction.

Previously identified tRNA substrates of RlmN include isodecoders for the following codons: GAA/GAG (glutamate), CAT/CAC (histidine), CAA/CAG (glutamine), CGT/CGC/CGA (arginine), and GAT/GAC (aspartate) in *E. coli* [82]. In *V. cholerae*, our results indicate that RlmN catalyzes the m²A37 modification in the isodecoders tRNA^Arg_ACG^ (decoding CGT and CGA codons), tRNA^Asp_GTC^ (decoding GAC and GAT codons), and tRNA^His_GUG^ (decoding CAC and CAT codons), but not in the other substrates observed in *E. coli*. Based on these targets, we selected the codons CGT/CGC/CGA (arginine), GAT/GAC (aspartate), and CAT/CAC (histidine) for analysis. Deletion of *rlmN* caused a modest but significant reduction in decoding efficiency for the CGT and CGA arginine codons (**Figure 7D**). Gene ontology (GO) analysis of genes enriched for these codons revealed categories associated with motility, translation, amino acid and nucleotide metabolism, and aromatic compound biosynthesis (**Figure 7E**).

*MiaA* catalyzes the isopentenylation of A37 in tRNAs decoding codons beginning with U [83], such as those for phenylalanine (TTT), leucine (TTA, TTG), serine (TCT, TCA, TCG, TCC), tyrosine (TAC, TAT), and cysteine (TGC, TGT). In *E. coli*, MiaA has been shown to influence the translation of *rpoS* and its regulator *iraP* by modulating leucine-UUX decoding [16, 84]. In *V. cholerae*, we observed consistent results: the absence of *miaA* led to reduced decoding efficiency for most U-starting codons (consistent with previous literature reviewed in [7]), with the notable exception of cysteine TGT (**Figure 7F**). The most pronounced effects were observed for tyrosine TAT and phenylalanine TTT. GO enrichment analysis of genes biased for these codons highlighted categories such as DNA repair and signal transduction (**Figure 7GH**), consistent with our previous findings [20].

Together, these results indicate that A37 modifications introduced by RlmN and MiaA fine-tune translation by modulating the efficiency of decoding specific, codon-biased transcripts, particularly those involved in stress response, regulation, and key metabolic pathways.

## Discussion

This study provides a comprehensive overview of the tRNA modification landscape in *Vibrio cholerae*, combining next-generation sequencing-based mapping approaches with targeted genetic deletions and phenotypic characterization. A key strength of our approach lies in the systematic validation of modification sites by monitoring their absence in strains lacking the corresponding modification enzymes, whenever feasible.

### Insights into the tRNA modification landscape of *Vibrio cholerae*

Although exhaustive tRNA modification maps exist for a limited number of model organisms as mentioned in the introduction, our work expands this repertoire by presenting the first detailed isodecoder-level map for *V. cholerae*, revealing several species-specific features, adding to the previously described C-to-Ψ editing by TrcP, acacp^3^U by AcpA and m^¹^A22 by TrmK [47]. Note that some tRNA modifications could not be detected with the sequencing approaches used here; Table 1 lists untested modifications with predicted homologous enzymes in *V. cholerae*, and certain modifications may be condition-specific and therefore absent under the growth conditions examined.

One peculiar feature of *V. cholerae* is the presence of *trmK*, which introduces the m¹A22 modification. While TrmK has been well characterized in Gram-positive bacteria like *Bacillus subtilis* [71], the vast majority of Gram-negative bacteria, including *E. coli,* lacks *trmK.* Its presence in *V. cholerae*, as well as in a handful of species of Gammaproteobacteria (*Vibrio* spp., *Shewanellea* spp.), suggests possible horizontal gene acquisition. The conservation of *trmK* in *V. cholerae* may reflect an evolutionary advantage in specific environmental niches or host interactions.

Comparative analysis of tRNA-modification enzymes across the *Vibrio* genus revealed a highly conserved repertoire within *Vibrio cholerae* strains, with all enzymes examined being present in 100 % of the genomes analyzed. Across *Vibrio* species, the canonical tRNA-modification enzymes are broadly conserved (**Figure S8**), with possibly varying regulatory control of modification levels [47]. For example, regulation of modification levels and peripheral steps of the queuosine (Q) pathway (salvage vs. de novo) can vary by lineage, with *V. cholerae* retaining the core Q-insertion machinery despite ecological tuning [85]. In contrast, a smaller accessory set, shows patchy, species-or strain-specific distribution (**Figure S8**), consistent with the highly flexible Vibrionaceae pan-genome [46, 86]: *trmK* is absent from 54 of the 106 species analyzed, *trhO* from 46, and *trcP* and *tapT* from 13 each. The absence patterns of *trcP* and *trmK* are consistent with their known taxonomic distribution, *trcP* having so far been reported as *Vibrio*-specific and *trmK* being generally associated with Gram-positive bacteria. These differences highlight lineage-specific remodeling of the tRNA-modification network in *Vibrio* and suggest potential functional diversification beyond the *V. cholerae* clade.

An interesting finding from this study is that, although the conserved Ψ55 modification is absent from most tRNAs in the *ΔtruB* mutant, both tRNA^Gln_TTG1^ and initiator tRNA^Met^ retain Ψ55 independently of TruB. This site is also unaffected by deletion of *truA*, *truC*, or *rluA*, and the enzyme responsible for installing Ψ55 in these tRNAs remains unidentified. Another difference from *E. coli* is the absence of the Gm18 modification in *V. cholerae* tRNAs, despite the presence of a predicted TrmH-family methyltransferase. This highlights that RNA modifications cannot always be reliably inferred from sequence homology alone and that further experimental work is needed to improve the accuracy of RNA modification prediction.

Furthermore, our data suggests interplay among Dus enzymes. In the absence of DusC, increased dihydrouridine formation by DusA and DusB is observed, potentially due to enhanced availability of the FMN cofactor otherwise utilized by DusC. Alternatively, DusC may compete with DusA/B for substrate binding, and its deletion could release this competitive inhibition, enabling increased activity by the remaining Dus enzymes. Interestingly, deletion of Dus enzymes, along with others, lead to decreased stop codon readthrough. Since D modification is known to introduce flexibility to tRNAs [87], one can speculate that its absence may increase rigidity, thereby limiting misdecoding. The idea that structural changes in the tRNA D-loop can impact decoding fidelity has been demonstrated through their influence on proofreading by the elongation factor, revealing a trade-off between fidelity and translation speed [88]. Such increase of accuracy in the absence of D can explain the decrease of stop codon readthrough observed in *dus* mutants.

### Insights on the effects of tRNA modifications on codon decoding and bacterial adaptation

Using a previously validated reporter system [20], we detect that loss of the U34-modifying enzymes TrhP or MnmE selectively impairs decoding of G-ending arginine and several leucine codons, while loss of the A37-modifying enzyme MiaA broadly reduces decoding of U-starting codons such as tyrosine TAT and phenylalanine TTT. Genes enriched for TrhP/MnmE-sensitive codons in *V. cholerae*, including key regulators such as the oxidative stress factor OxyR, are overrepresented in pathways linked to protein complex assembly and stress responses, whereas MiaA-sensitive codons are enriched in DNA repair, supporting a model in which codon usage and tRNA modification together tune translation to regulate stress adaptation in *V. cholerae*.

Together, these findings highlight both the conserved and divergent features of tRNA modification biology in *V. cholerae* and lay the groundwork for exploring the functional consequences of these modifications in stress responses, environmental adaptation, and pathogenicity.

### Species-specific differences in tRNA isodecoder repertoires shape the impact of modifications

An important factor that may influence the species-specific impact of tRNA modifications is the number and diversity of tRNA isodecoder genes encoded in the genome. Comparative analysis between *E. coli* and *V. cholerae* reveals substantial differences in isodecoder copy number and repertoire (**Figure 8**). Notably, several tRNA genes present in *E. coli* are absent in *V. cholerae*, including those decoding glycine (GGG), arginine (AGG), glutamine (CAG), proline (CCG), and serine (UCG). Interestingly, despite similar codon usage frequencies for glycine GGG and the rare arginine AGG codons in both species, the usage of glutamine CAG and proline CCG codons is significantly lower in *V. cholerae* (35% and 27%, respectively) compared to *E. coli* (65% and 53%).

**Figure 8.**
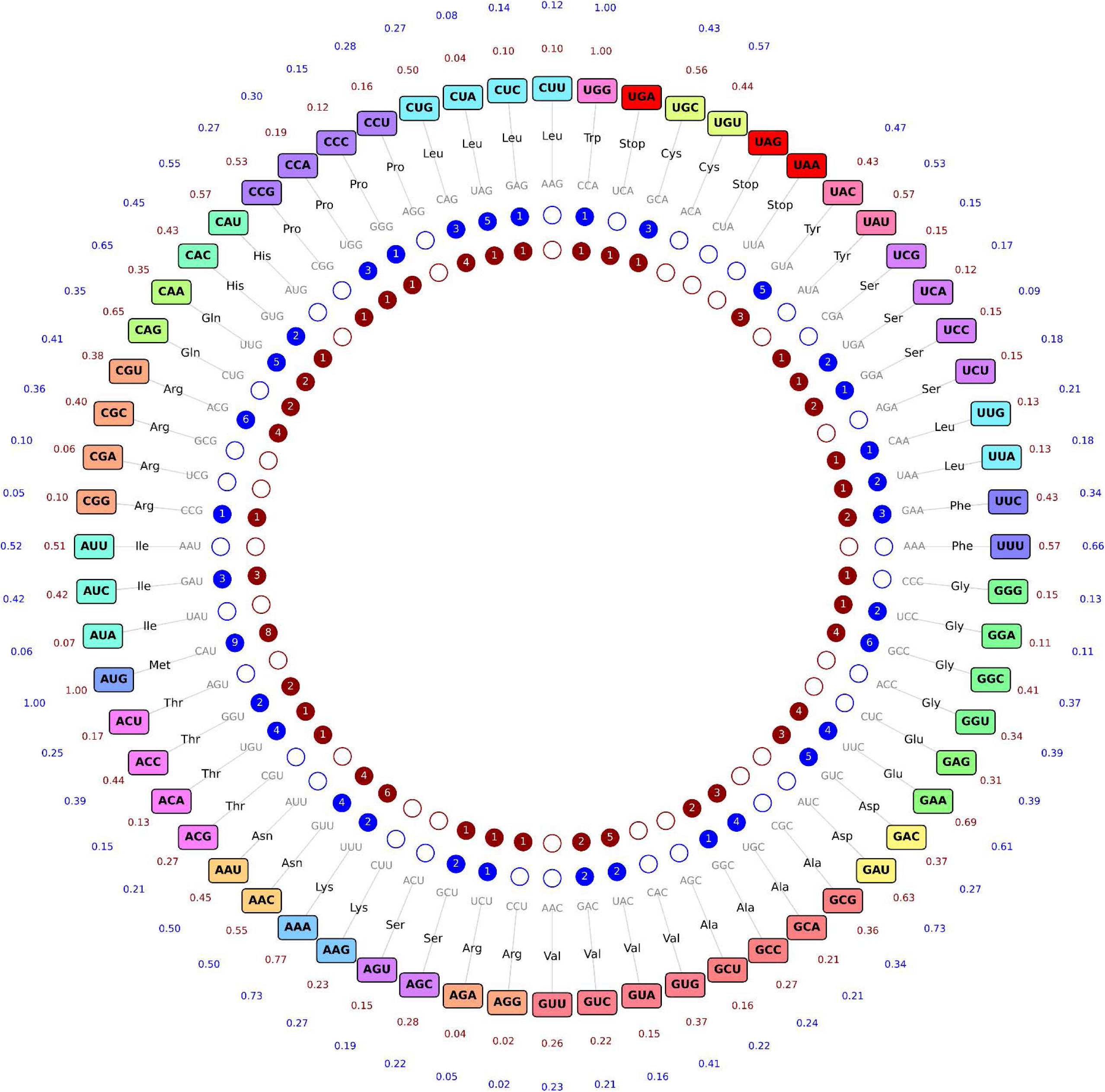
Comparison of the number of isodecoders and codon usage between *V. cholerae* and *E. coli*. Codons are arranged around a circle and color-coded according to their corresponding amino acid. For each codon, the relative usage frequency is displayed outside the circle in blue for *V. cholerae* and brown for *E. coli*. The reverse-complement anticodon is positioned opposite, and the number of tRNA isoacceptors is shown in inner dots in blue for *V. cholera*e and brown for *E. coli*.

Isodecoder copy numbers also vary substantially between the two species. For example, tRNA^Leu_CUA^ is present in five copies in *V. cholerae* versus one in *E. coli*; similarly, tRNA^Thr_ACA^ (4 vs. 1) and tRNA^Cys_UGC^ (3 vs. 1) exhibit notable differences. These disparities suggest that species-specific tRNA gene content can influence both the reliance on and functional consequences of particular modifications.

For instance, in *V. cholerae*, where only a single tRNA^Gln^ isodecoder (UUG anticodon) exists, modifications in the T-and D-arms may be critical for decoding both glutamine codons efficiently. In contrast, *E. coli*, which possesses two distinct tRNA^Gln^ isodecoders (UUG & CUG anticodons), may be less dependent on such modifications for decoding flexibility. Similarly, the m¹G37 modification (catalyzed by TrmD) in tRNA^Pro^ with UGG/GGG anticodons could be essential in *V. cholerae* for decoding proline CCG codons, given the absence of redundant isodecoders. However, TrmD is essential, and thus its deletion could not be tested directly.

Altogether, this study adds a new layer of understanding to the diversity of tRNA modifications and their modifiers in *V. cholerae*. These findings emphasize the importance of expanding such analyses to additional species, particularly those with unique ecological niches or stress responses. Exploring species-specific tRNA repertoires and modification patterns can reveal novel mechanisms of translational control and adaptation, as well as previously uncharacterized modifications.

## Materials and methods

### Bacterial strain constructions

All strains used in this study are derivatives of *Vibrio cholerae* O1 biovar El Tor N16961 hapR⁺, and were constructed via allelic exchange. Mutant strains were generated using the *ΔlacZ* strain (K329) as the parental background. Detailed information on strain construction and primer sequences is provided in **Table S1.** Cultures were grown at 37°C in Mueller–Hinton (MH) medium, both in solid and liquid form, unless indicated otherwise. The *mnmE*, *trmF*, *trmA*, *trmB*, *rlmN*, *tgt*, *dusA*, *dusB*, *truA*, *truB*, *truC*, *trmK*, and *miaB* deletion mutants were constructed as previously described [19]. The *trmJ*, *trmL*, *ttcA*, *trcP*, *trhP*, *thiI*, *dusC*, *truD*, *miaA*, *rluA*, *tapT*, and *trmO* mutants were generated by transformation or co-transformation [89], as indicated in Table S1.

For standard natural transformation, the PCR product was generated by overlap extension to replace the target gene with a kanamycin resistance cassette flanked by ∼300–500 bp of homology to the chromosomal regions immediately upstream and downstream of the gene.

For co-transformation, two PCR products were co-introduced: one unmarked deletion fragment generated by overlap extension PCR, consisting of ∼3 kb of upstream and downstream homology flanking the gene to be deleted, and one PCR product carrying a chloramphenicol resistance cassette flanked by ∼500 bp of homology targeting a neutral chromosomal locus: the intergenic region between *dsbB* (VC1902) and *ftsK* (VC1903). Selection was performed on the cassette, while the unmarked deletion was validated by PCR screening.

Natural transformation was performed as follows. *V. cholerae* cells (*ΔlacZ* strain K329) were grown in LB to an OD_₆₀₀_ of 0.8, pelleted by centrifugation, and resuspended in 1 mL of Instant Ocean. A 100 µL aliquot was used to inoculate 2 mL Eppendorf tubes containing 150 µL of chitin slurry and 750 µL of Instant Ocean. Cultures were incubated statically at 30°C for 16 hours to induce competence. After incubation, 550 µL of the supernatant was removed, and DNA was added: 100 ng of PCR product for transformation, or a mixture of 100 ng of the chloramphenicol cassette and 3 µg of the unmarked deletion fragment for co-transformation. Tubes were incubated for another 16 hours at 30°C. After this second incubation, cultures were vortexed vigorously, 500 µL of LB was added, and cells were incubated for 2 hours at 37°C with shaking. Finally, 100 µL of the culture was plated on LB agar containing either kanamycin (25 µg/mL) or chloramphenicol (5 µg/mL), depending on the selection used.

### tRNA enriched RNA extraction

*V. cholerae* overnight cultures were diluted 1:1000 in Mueller–Hinton (MH) medium and incubated aerobically at 37°C with shaking at 180 rpm until reaching an OD₆₀₀ of 0.25. For H_2_O_2_ treated samples: cultures were then treated or not with 1mM H_2_O_2_ for 10min. Total RNA enriched in tRNAs was extracted at room temperature using TRIzol reagent, following a previously described protocol [20]. Genomic DNA was removed using the TURBO DNA-free Kit (Ambion), according to the manufacturer’s instructions. RNA concentration was determined with a NanoDrop 2000c spectrophotometer (Thermo Fisher Scientific), and the quality of tRNA-enriched fractions was assessed by capillary electrophoresis using a RNA 6000 Pico chip on a Bioanalyzer 2100 (Agilent Technologies).

### tRNA reference sequence for *Vibrio cholerae* O1_biovar_El_Tor_str_N16961

Comprehensive analysis of tRNA modifications is a challenging task, even for simple model organisms, like bacteria, which have only a limited number of tRNA-encoding genes. We first constructed a non-redundant tRNA reference for *Vibrio cholerae* O1 biovar El Tor str N16961. *Vibrio cholerae* O1_biovar_El_Tor_str_N16961 [90] genome encodes 55 tRNA genes, some of them are either identical or show substantial sequence similarity (1-5 nucleotide replacements/mutations). In order to construct non-redundant tRNA reference, we first identified identical tRNA sequences and collapsed them into unique tRNA isoacceptor, and further collapsed closely related tRNA species in a single sequence. Detailed procedure was described previously [91]. This “collapsed” reference sequence contained 38 tRNA sequences, species showing <12 nt substitutions were collapsed in one sequence entry. Resulting tRNA sequences were further modified by adding 3’-CCA sequence (if absent in the encoded genomic sequence) and by changing A34 (if any) to G34, since A at the tRNA wobble position is systematically converted to Inosine(I34) which base-pairs with C during the RT-step of library preparation. This amended collection of *V. cholerae* tRNAs was further used for analysis of NGS RNA modification mapping data.

### Mapping of tRNA modifications in *V. cholerae* by deep sequencing-based methods

The tRNA map was created in BioRender. Baharoglu, Z. (2025) https://BioRender.com/i9kiyrr.

Each detection experiment described below was performed both for wild-type *V. cholerae* and the corresponding deletion mutants. This approach allowed us to unambiguously assign specific modification signals to their respective enzymes by comparing their presence in wild-type strains with their absence in the relevant gene deletion backgrounds. Note that the detection of queuosine modification of tRNA^Asp^, tRNA^Tyr^, tRNA^Asn^ and tRNA^His^, made by the Tgt enzyme, was previously described in our previous study of *V. cholerae* tRNA modifications [20]. Deep sequencing-based approaches for mapping 2′-O-methylated (Nm) and pseudouridylated (ψ) residues, respectively RiboMethSeq [32, 34] and HydraPsiSeq [35, 66, 92], differ in the chemical cleavage strategies they employ, but are conceptually similar in the type of signal they detect. Both methods rely on the relative resistance of modified RNA residues to specific chemical cleavage, compared to their unmodified counterparts. AlkAnilineSeq [33, 36] and RiboMethSeq [34] have been previously described in details.

AlkAnilineSeq relies on inducing labile N-glycosidic bonds via alkaline hydrolysis, followed by aniline-mediated cleavage at RNA abasic sites. This process releases RNA fragments with terminal 5’-phosphates, enabling specific ligation of sequencing adapters. Due to this selective ligation strategy, AlkAnilineSeq produces high signal-to-noise ratios and minimal background from unmodified RNA. Among the detectable modifications, m⁷G methylation, typically stoichiometric, yields strong signals, whereas substoichiometric modifications like D are also detectable, albeit with lower signal intensity. m⁷G and D residues are universally present in bacterial tRNAs, allowing for their precise and reproducible mapping using this method.

In RiboMethSeq, alkaline hydrolysis is used to randomly fragment RNA, and the 3′-phosphodiester bond adjacent to Nm residues is more resistant to cleavage. In contrast, HydraPsiSeq utilizes hydrazine treatment to selectively cleave unmodified uridines, while adenine, cytosine, guanine, and ψ residues remain intact, resulting in a protection signal at ψ sites. Notably, these protocols are based on negative detection; that is, they infer the presence of modifications from protection against cleavage, rather than through direct modification-specific signatures. Consequently, both methods can be more prone to false positives compared to positive detection approaches such as BS-Seq or AlkAnilineSeq. To increase specificity, the use of deletion strains lacking individual modifying enzymes provides a powerful strategy for unambiguous assignment of modification sites to their corresponding enzymes. This is especially important given that both Nm methyltransferases and pseudouridine synthases belong to large and functionally diverse enzyme families, and their substrate specificities often cannot be accurately predicted based on sequence similarity to known homologs. Therefore, complementary mapping approaches for Nm and ψ modifications are essential for orthogonal validation and for increasing confidence in site-specific modification assignment.

### Bisulfite sequencing (BS-Seq)

tRNAs were subjected to bisulfite treatment according to published protocols [69, 93, 94]. BS-Seq analysis of m^5^C locations was performed using alignments of the sequencing reads to a C to U converted reference sequence, by determining the residual C non-deamination. The proportion of residual non-deaminated C (UtoC score) corresponds to the molar ratio of m^5^C at a given location. The UtoC score was calculated from full mpileup format, as described above for BID-Seq analysis.

**GLORI** is based on deamination, like BS-seq. The GLORI protocol includes a glyoxal treatment that serves as a ’protection’ step, selectively masking guanosine residues from nitrite (NO₂⁻)-induced deamination. This is followed by deprotection and reverse transcription (RT) to convert RNA into cDNA. The conditions used were as described [95]. Briefly 100 ng of RNA were subjected to glyoxal treatment for 30 min at 50°C followed by boric acid treatment for 30 min at 50°C. The protected RNA is then subjected to nitrite-mediated deamination followed by RNA deprotection in a buffer containing 500 mM TEAA and formamide. RNA was then precipitated, end-repaired and purified before being subjected to library preparation with NEBNext small RNA library following manufacturer’s protocol. The quality and quantity of each library were assessed using a high-sensitivity DNA Chip on a Bioanalyzer 2100 and a Qubit 2.0 fluorometer. High-throughput sequencing of the multiplexed libraries was performed on an Illumina NextSeq 2000 instrument in a 75 nt single-end read mode or 2×50 nt paired-end mode. Raw sequencing reads were inspected with FastQC and adapter sequence was removed by trimmomatic v0.3931. Alignment to the A->G converted tRNA/rRNA reference sequence was done by Bowtie2. v2.4.232 with slightly relaxed alignment stringency, allowing to retain 1 nt mismatched reads. Further analysis was done by samtools mpileup utility and counting the mismatch profile at every position in the reference. Mismatches GtoA correspond to deamination-resistant modified residues (m6A and all other N6-modified A, m1A was also detected as GtoA signal). The value of GLORI GtoA score corresponds to the molar ratio of the modified A.

### BID-Seq

To assess the presence/absence of pseudouridine in tRNAs, a derived version of BID-Seq protocol was performed [96]. Briefly, 100 ng of RNA were subjected to RNA fragmentation followed by bisulfite treatment and desulphonation as described [97]. Treated RNAs were end-repaired, purified, and subjected to NEBNext small RNA library following manufacturer’s protocol. The quality and quantity of each library were assessed using a high-sensitivity DNA Chip on a Bioanalyzer 2100 and a Qubit 2.0 fluorometer. High-throughput sequencing of the multiplexed libraries was performed on an Illumina NextSeq 2000 instrument in a 75 nt single-end read mode. Raw sequencing reads were inspected with FastQC and adapter sequence was removed by trimmomatic v0.3931. Alignment to the reference sequence was done by Bowtie2. v2.4.232 with “relaxed” alignment stringency, allowing to retain 1–3 nt gapped reads. Further analysis was done by samtools mpileup utility and counting the deletion score at every position in the reference. Deletions/jumps correspond to bisulfite-modified residues. Background value of the deletion score (mock-treated RNA sample) was subtracted.

### RNA-seq and RNA editing analysis

We used a modified version of our RNA editing detection pipeline published previously [98]. Specifically, we used CLC Genomics Workbench for all analysis steps (described below).

RNA-seq reads were first trimmed according to length and quality scores to ensure high quality of the reads by using the following parameters: Trim using quality scores = Yes; Quality limit = 0.01; Trim ambiguous nucleotides = Yes; Maximum number of ambiguities = 1; Automatic read-through adapter trimming = Yes; Minimum length = 50; Maximum length = 150; Remove 5’ terminal nucleotides = No; Remove 3’ terminal nucleotides = No; Remove on first read = Yes; Remove on second read (for paired reads) = Yes; Trim to a fixed length = No; Trim end = Trim from 3’-end; Discard short reads = Yes; Discard long reads = No; Save discarded sequences = No.

Next, RNA-seq reads were mapped to the *V. cholerae* reference genome (chromosomes NC_002505 and NC_002506) with the following parameters: Masking mode = No masking; Match score = 1; Mismatch cost = 2; Cost of insertions and deletions = Linear gap cost; Insertion cost = 3; Deletion cost = 3; Length fraction = 0.8; Similarity fraction = 0.8; Global alignment = No; Non-specific match handling = Map randomly. Next, initial variant calling was performed using the following parameters: Ignore positions with coverage above = 10,000,000; Restrict calling to target regions = Not set; Ignore non-specific matches = No; Minimum coverage = 1; Minimum count = 1; Minimum frequency (%) = 0.1; Base quality filter = Yes; Neighborhood radius = 5; Minimum central quality = 30; Minimum neighborhood quality = 30; Read direction filter = No; Relative read direction filter = No; Read position filter = No.

After the initial variant calling, additional filtering was applied to identify editing events transcribed from the positive strand. All filtered variants were required to match all the following criterions: Criteria = ’Type contains SNV’; Criteria = ’Reference allele contains No’; Criteria = ’Frequency >= 1’; Criteria = ’# unique start positions >= 3’; Criteria = ’# unique end positions >= 3’; Criteria = ’Reverse read count >= 3’; Criteria = ’Reverse read coverage >= 10’; Criteria = ’Reference contains A and the variant contains G.

After the initial variant calling, additional filtering was applied to identify editing events transcribed from the negative strand. All filtered variants were required to match all the following criterions: Criteria = ’Type contains SNV’; Criteria = ’Reference allele contains No’; Criteria = ’Frequency >= 1’; Criteria = ’# unique start positions >= 3’; Criteria = ’# unique end positions >= 3’; Criteria = ’Forward read count >= 3’; Criteria = Forward read coverage >= 10’; Criteria = ’Reference contains T and the variant contains C.

All variants were combined into a single list used as a reference to extract all variants’ information from each sample’s initial variant’ files, allowing us to identify the state of editing in sites and samples filtered out due to our filters.

Next, we filtered for variants shared between at least two (out of three) biological replicates. Next, we extracted the status of the identified shared variants from the RNA mapping step to ensure we did not miss variants because of our filters and to ensure that when a variant is absent in our initial analysis, it is not because its transcript was not sequenced.

Finally, we focused only on sites in annotated tRNA genes. Importantly, only sites in transcript tRNA^Arg2^ were discovered.

### Competition experiments

Overnight cultures from single colonies of *lacZ⁺* mutant and *lacZ⁻* wild-type strains (or vice versa) were washed with phosphate-buffered saline (PBS) and mixed in a 1:1 ratio (500 µl each). To assess the initial T_₀_ ratio, 100 µl of the mixture were serially diluted and plated on MH agar containing X-gal (40 µg/ml). Simultaneously, 10 µl of the mixture were inoculated into 2 ml of MH medium (approximately 5 × 10^5^ cells/ml), either alone or supplemented with sub-inhibitory tobramycin (0.6 µg/ml). Cultures were incubated at 37°C with agitation for 20 hours (∼9 generations). Following incubation, cultures were diluted and plated on MH-X-gal plates. After overnight growth at 37°C, blue (*lacZ⁺*) and white (*lacZ⁻*) colonies were counted. The competitive index was calculated by dividing the number of *lacZ⁺* CFUs by *lacZ⁻* CFUs and normalizing to the initial T₀ ratio.

**Construction of *gfp* reporters with codon stretches [20]:** As a positive control, we used *gfpmut3*, a stable variant of GFP expressed under the control of the *V. cholerae PgyrA* promoter. The full reporter sequence is provided below, with the **–35** and **–10** promoter elements, as well as the **ATG** start and **TAA** stop codons, indicated in **bold and underlined**. The site for insertion of the test stretch is shown in *italics*.

TGACTTGGCGCTCAATCTTGTAGTGAGC**<u>TTCGTT</u>**TCAGTAAGAATTT**<u>GGGTATACC</u>**GATCAAACTATAGAGGGATA ATGGCTCT**<u>ATG</u>***(±stretches)*CGTAAAGGAGAAGAACTTTTCACTGGAGTTGTCCCAATTCTTGTTGAATTAGATGGTGATGTTAATGGGCACAAATTTTCTGTCAGTGGAGAGGGTGAAGGTGATGCAACATACGGAAAACTTACCCTTAAATTTATTTGCACTACTGGAAAACTACCTGTTCCATGGCCAACACTTGTCACTACTTTCGGTTATGGTGTTCAATGCTTTGCGAGATACCCAGATCATATGAAACAGCATGACTTTTTCAAGAGTGCCATGCCCGAAGGTTATGTACAGGAAAGAACTATATTTTTCAAAGATGACGGGAACTACAAGACACGTGCTGAAGTCAAGTTTGAAGGTGATACCCTTGTTAATAGAATCGAGTTAAAAGGTATTGATTTTAAAGAAGATGGAAACATTCTTGGACACAAATTGGAATACAACTATAACTCACACAATGTATACATCATGGCAGACAAACAAAAGAATGGAATCAAAGTTAACTTCAAAATTAGACACAACATTGAAGATGGAAGCGTTCAACTAGCAGACCATTATCAACAAAATACTCCAATTGGCGATGGCCCTGTCCTTTTACCAGACAACCATTACCTGTCCACACAATCTGCCCTTTCGAAAGATCCCAACGAAAAGAGAGACCACATGGTCCTTCTTGAGTTT GTAACAGCTGCTGGGATTACACATGGCATGGATGAACTATACAAA**<u>TAA</u>**

To assess amino acid incorporation efficiency at specific codons, three tandem repeats of the target codon were inserted directly downstream of the *gfp* gene’s ATG start codon. Double-stranded DNA fragments were synthesized as eBlocks (Integrated DNA Technologies, IDT) and cloned into the pTOPO-Blunt vector, which carries a kanamycin resistance marker, according to the manufacturer’s instructions.

### Quantification of *gfp* fusion expression by fluorescent flow cytometry

After Miniprep plasmid purification, constructs were transformed into the target strains. Flow cytometry was performed on overnight cultures as previously described[20], with each experiment repeated at least six times. For each replicate, 50,000 to 100,000 events were recorded using a Miltenyi MACSQuant flow cytometer. Mean fluorescence per cell was measured in the FITC channel for each reporter in both wild-type and mutant backgrounds.

To assess codon decoding efficiency, the ΔΔfluorescence ratio was calculated as follows: fluorescence of the codon-stretch reporter in the mutant strain was divided by the mean fluorescence of the same reporter in the wild-type strain (Δfluorescence_codon X_). This value was then normalized to the Δfluorescence of a control reporter lacking codon inserts (Δfluorescence_wt gfp_). This two-step normalization accounts for baseline differences in GFP translation between wild-type and mutant strains, enabling a more accurate comparison of codon-dependent decoding efficiency.

**Stop codon readthrough** was performed as described [20, 75].

### Gene Ontology (GO) enrichment analysis

GO enrichment analysis was performed using the tools available at geneontology.org. Specifically, Fisher’s exact test was used to determine whether certain gene groups in the input list were significantly over-or underrepresented compared to a reference set. The analysis utilized the GO Biological Process Complete annotation dataset. Input gene lists corresponded to genes exhibiting codon usage bias for specific codons of interest, as defined in previously published datasets [20] (https://zenodo.org/records/6875293). The reference set included all 3,782 annotated *V. cholerae* genes. The analysis was conducted using the PANTHER Overrepresentation Test (version released 2022-07-12) with GO Ontology database version 2022-03-22 (DOI: 10.5281/zenodo.6399963).

### Quantification and statistical analysis

For comparisons between 2 groups, first an F-test was performed in order to determine whether variances are equal or different between comparisons. For comparisons with equal variance, Student’s t-test was used. For multiple comparisons, we used ANOVA to determine the statistical differences (*p*-value) between groups. **** means p<0.0001, *** means p<0.001, ** means p<0.01, * means p<0.05.

Comparison of the number of isodecoders and codon usage between *V. cholerae* and *E. coli*. Coding sequences (CDS) and tRNA genes were extracted from the GenBank files of *V. cholerae* N16961 chromosomes I and II and *E. coli* MG1655 using custom Python scripts. In a first step, all annotated tRNA genes were parsed to retrieve their product name, anticodon sequence, locus tag, and genomic position. For isoacceptors associated with the same amino acid, the corresponding sequences were aligned, and potential point mutations were identified by comparing them to a reference sequence. In a second step, all protein-coding sequences were used to compute codon usage frequencies. Codons were counted across all CDS and normalized within each amino acid group to yield relative usage values. Finally, a circular visualization was generated to integrate the codon usage data with tRNA information. The figure was created using Matplotlib.

### Distribution and sequence conservation of tRNA-modification enzymes across the *Vibrio* genus

Protein sequences of tRNA-modification enzymes from *Vibrio cholerae* O1 biovar El Tor strain N16961 (GenBank accessions NC_002505 and NC_002506) were used as queries to assess the distribution of homologs across the genus *Vibrio*. The complete *Vibrio* proteome (taxonomic identifier 662) was downloaded from UniProtKB in October 2025 (1,154,790 protein entries) and used as the target database. Sequence similarity searches were performed using DIAMOND BLASTp (v2.1.10) with an E-value cutoff of 1e−5, a minimum sequence identity of 30%, and a minimum query coverage of 60%. For each *V. cholerae* reference protein, up to 20,000 target sequences were retained, and the best hit per species was selected based on the highest percentage identity. Taxonomic information (species name and TaxID) was extracted from UniProt headers (fields OS= and OX=) and processed with custom Python scripts (v3.10) using the pandas and numpy libraries. Heatmaps were generated with matplotlib, where cell colors represent the maximum pairwise identity between each *V. cholerae* protein and its homolog in the corresponding *Vibrio* species. A condensed matrix was constructed by selecting one representative strain per species (based on its TaxID, preferentially reference or type strains), and this dataset was used to visualize the phylogenetic distribution of tRNA-modification enzymes across the genus. Only one representative genome per *Vibrio* species is shown. The resulting percent identity matrix was visualized as a heatmap, with color intensity representing sequence identity (yellow = 100 %, purple = 0 % / absence). Genes with no detectable hit (no significant BLAST alignment) were considered absent.

## Acknowledgments

We thank Valérie de Crécy-Lagard for valuable discussions and for generously sharing her knowledge. This research was funded, in whole or in part, by the Institut Pasteur (LH, ZB, DM, LF, EK), the Centre National de la Recherche Scientifique (CNRS-UMR 3525) (LH, ZB, DM, LF, EK), ANR ModRNAntibio (ANR-21-CE35-0012) (ZB, LH), ANR EpiRNA (ANR-24-CE12-7224) (ZB, LF) and the Fondation pour la Recherche Médicale (FRM EQU202103012569) (ZB, DM, LF). LH was funded by Institut Pasteur Roux-Cantarini fellowship. We thank Aurore Treffkorn-Maurau for help with competition experiments. For the purpose of open access, the authors have applied a CC-BY public copyright licence to any Author Manuscript version arising from this submission. The funders had no role in study design, data collection and analysis, decision to publish, or preparation of the manuscript. The authors have declared that no competing interests exist.

## Data availability

Sequencing data is publicly available through the ENA public database with accession number PRJEB93809. https://www.ebi.ac.uk/ena/browser/view/PRJEB93809

## Supplementary figures

**Figure S1: m²A signal detection with GLORI, attributed to RlmN.** Histograms show the A-to-G deamination rate (y-axis) at the indicated nucleotide positions (x-axis). A→G substitutions indicate unmodified adenosines, whereas retention of A signal (GtoA score, see legend to the Figure 1) denotes the presence of a modification. Note that the RlmN-dependent signal was weak and detected only in the three tRNAs shown.

**Figure S2. Mapping of pseudouridines (**Ψ**) and T54 by HydraPsiSeq. AB:** Detection of Ψ32 made by RluA (A) and C to U conversion by TrcP, followed by conversion to Ψ32 (B). HydraPsiSeq uses hydrazine to cleave unmodified uridines, with Ψ residues resisting cleavage and producing a protection signal. With BID-Seq, Ψ positions show up as characteristic single-base deletions (or stops) compared to the reference. **C**: Heatmap of m^5^U (T54) modifications detected by HydraPsiSeqin WT, and disappearance in *ΔtrmA.* Blue indicates absence of modification while yellow means presence **D:** Detection of k^2^C34 in WT as cleavage at C34 residue in tRNA^Ile_CAT^.

**Figure S3. A**. **Bisulfite-sequencing** of tRNAs from WT strains under non-treated (NT) and H_2_O_2_ treated conditions, showing the absence of m^5^C modification in tRNA^Tyr^. s^4^U at positions 8 and 9 are detected. **B**. **Analysis of RT-signatures for mapping of RT-mismatching and RT-arresting tRNA modifications.** Detection of s^4^U at positions 8 and 9 in WT and the absence of the signal in the deletion mutant for *ΔthiI*. acp^3^U47 detected in WT but not in *ΔtapT*. ms^2^i^6^A37 signal detected in WT but not in *ΔmiaB*.

**Figure S4: Phylogenetic distribution of TrmK (KO: K06967) according to the AnnoTree database.** AnnoTree visualizes annotations across a large phylogenetic tree, here comprising 80,000 bacterial genomes (one genome per species). *trmK* was found in 17,363 genomes. Absence (grey) or presence (colored) of an annotated *trmK* homolog is shown for each genome. The position of the Vibrionaceae family is indicated with a red arrow. Each phylum name is color-coded.

**Figure S5. Impact of RNA modification gene deletions on fitness during growth in sub-MIC tobramycin**.

**A.** Inosine frequency on tRNA^Arg^ detected in RNA-seq performed on *V. cholerae* without and with subinhibitory concentration of tobramycin (TOB, 0.4 µg/ml). For statistical analysis, student’s t-test was used. * means p<0.05. **B.** *In vitro* competition experiments of *V. cholerae* WT and mutant strains in the absence or presence of tobramycin at sub-MICs (50% of the MIC): 0.6 μg/ml. *y*-axis: log_2_ of competitive index value of the tested strain against the WT, calculated as described in the methods. Values shown for WT are competitions between 2 isogenic *lacZ+* vs *lacZ-* WT strains. A competitive index of 1 indicates equal growth of both strains. NT: no antibiotic treatment. For multiple comparisons, we used one-way ANOVA **** means *P*< .0001, *** means *P*< .001, ** means *P*< .01, and * means *P*< .05. Only significant *P*-values are represented. Number of biological replicates for each experiment: n=3.

**Figure S6**: **Stop codon readthrough for tRNA modification gene deletion mutants.** Black: wild-type (WT). Red: *Δmutant*. NT: no treatment. TOB: growth in the presence of tobramycin at 20% of the MIC. Reporters described in <u>Fabret and Namy, 2021</u>, and Fruchard et al 2025. Y-axis represents stop codon readthrough*1000. Number of biological replicates: between 3 and 6. For multiple comparisons, we used one-way ANOVA. **** means p<0.0001, *** means p<0.001, ** means p<0.01, * means p<0.05. Venn diagrams showing overlap of tRNA-modification enzyme deletion mutants that (orange) increase or (blue) decrease stop-codon readthrough across UAA, UAG, and UGA. Small circles indicate the most shared tRNA isodecoders among enzymes in each group; for instance, UAA readthrough is reduced in *thiI* and *trmK* mutants, both targeting tRNA^Tyr_GTA^.

**Figure S7:** Heatmap for arginine and leucine codon usage for transcription factors identified as biased for Arg AGG/CGG and Leu TTA/TTG/CTA codons over the other Arg and Leu codons. Standardized codon usage bias is shown for the codons of interest. In the plots, **red** indicates a positive codon usage bias (the codon occurs more frequently than expected relative to the genome-wide mean), whereas **blue** indicates a negative codon usage bias (the codon is underrepresented compared with the genomic average).

**Figure S8. Distribution and sequence conservation of tRNA-modification enzymes across the *Vibrio* genus.** Heatmap showing the presence and sequence identity of tRNA-modification enzymes (rows) across *Vibrio* species (columns). Color intensity represents amino acid sequence identity relative to the *Vibrio cholerae* N16961 reference, with yellow indicating 100 % identity and shades toward purple indicating lower identity. Purple squares denote complete absence of the corresponding gene. All *V. cholerae* strains show full presence of the enzymes

**Table S1: Strains and primers**

