## Supplementary figures and images for "The tRNA epitranscriptomic landscape and RNA modification enzymes in *Vibrio cholerae*"

### Figure S

GLORI, $m^2A37$ by RlmN

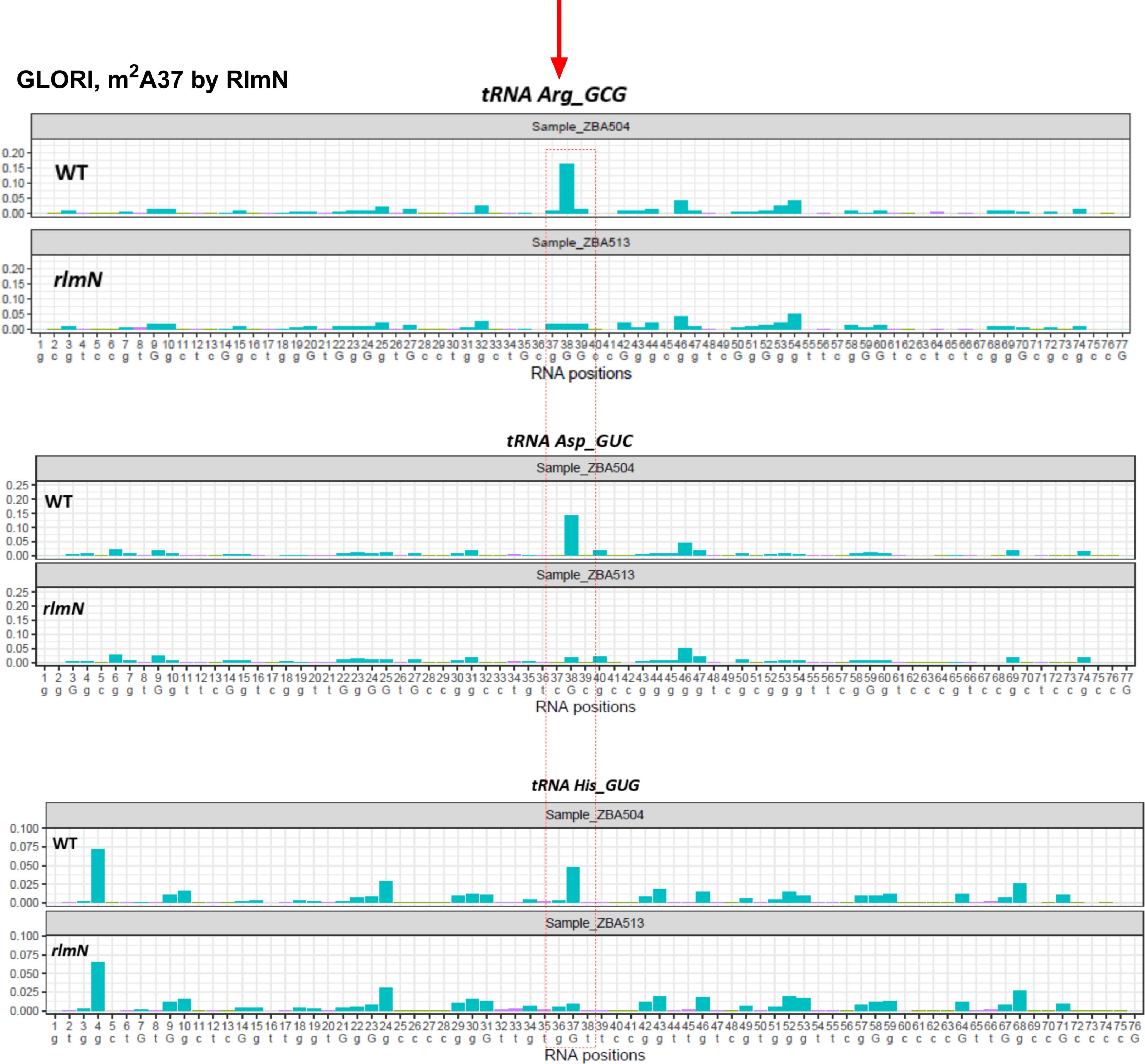

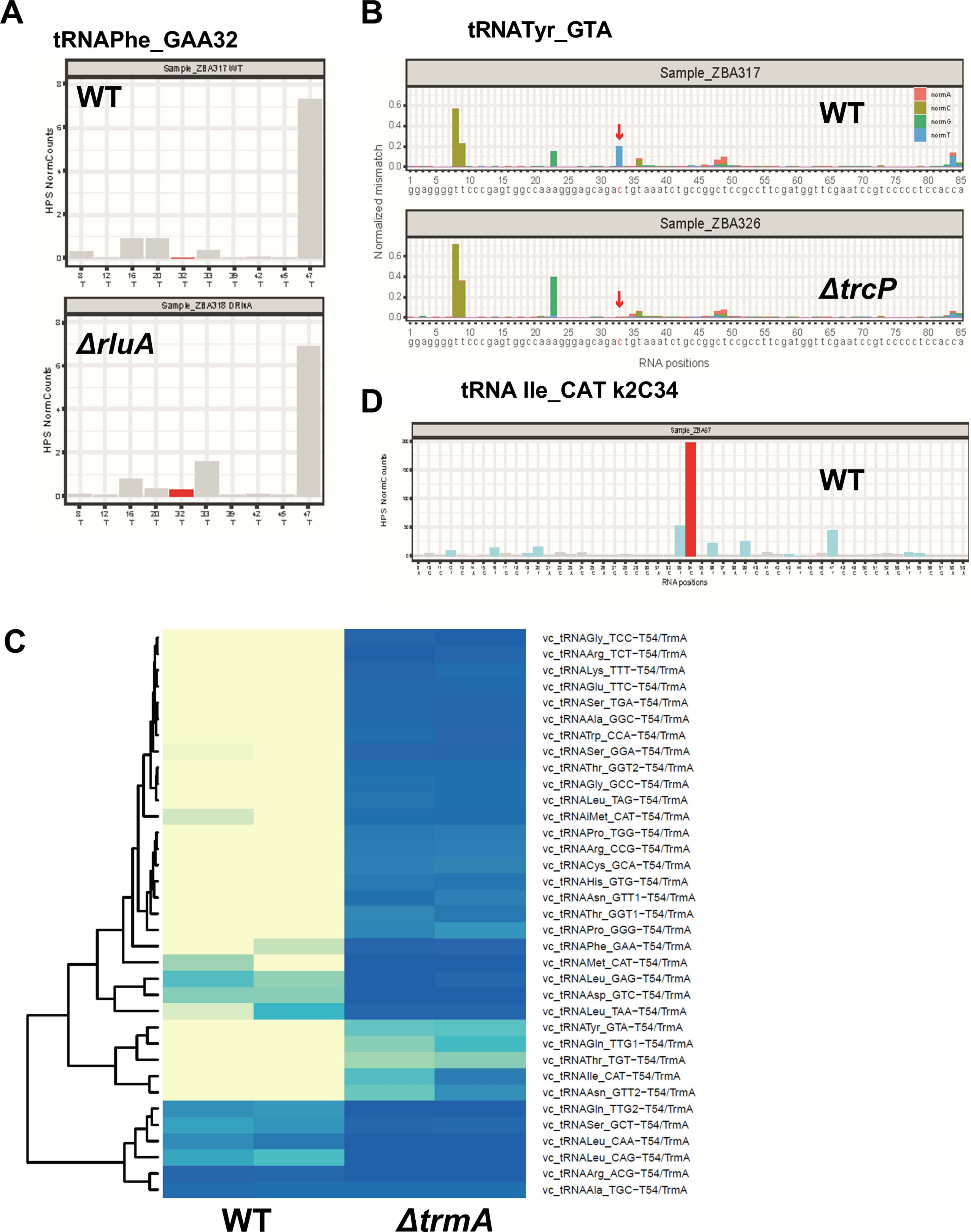

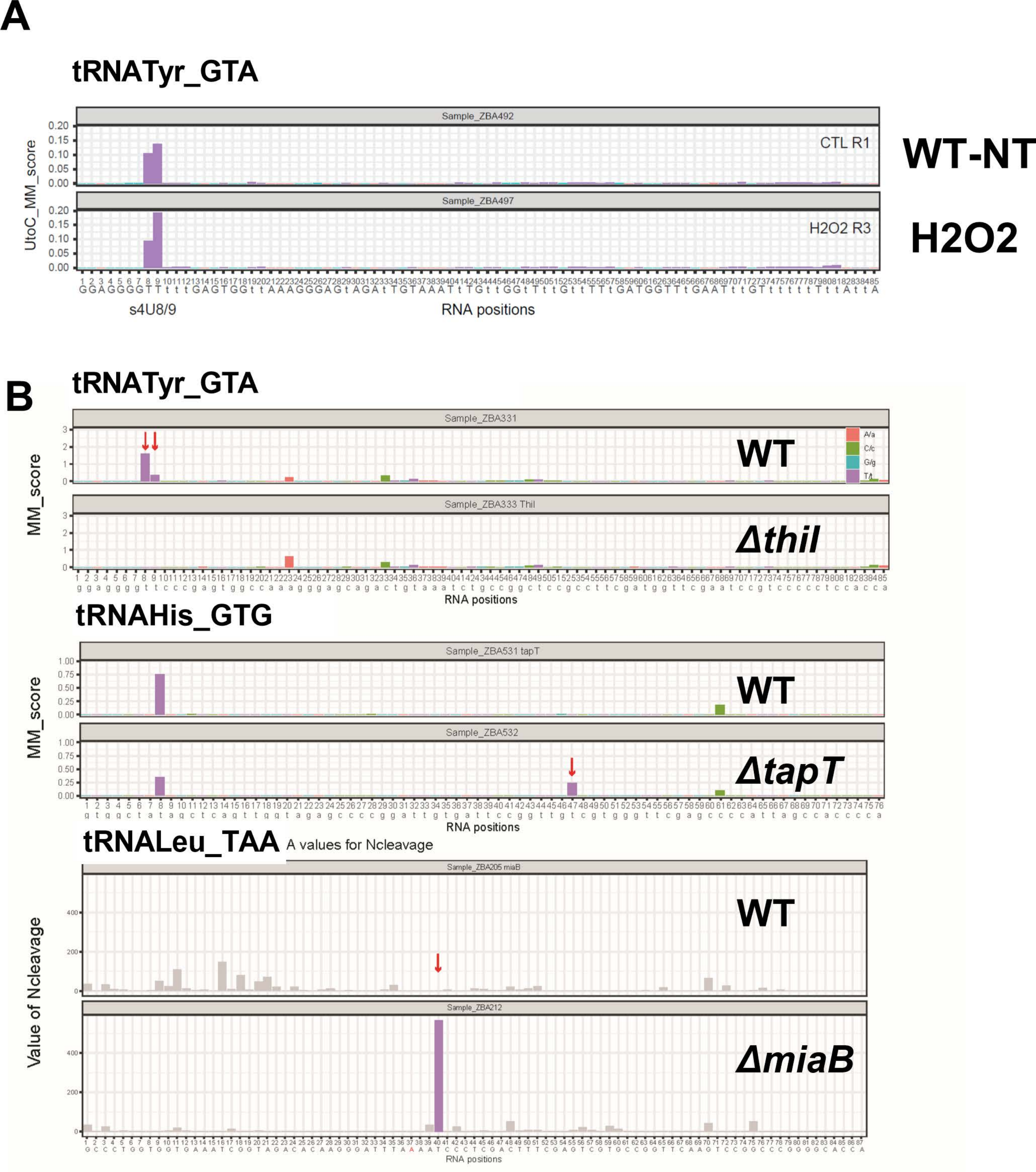

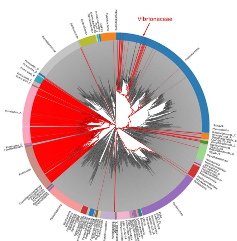

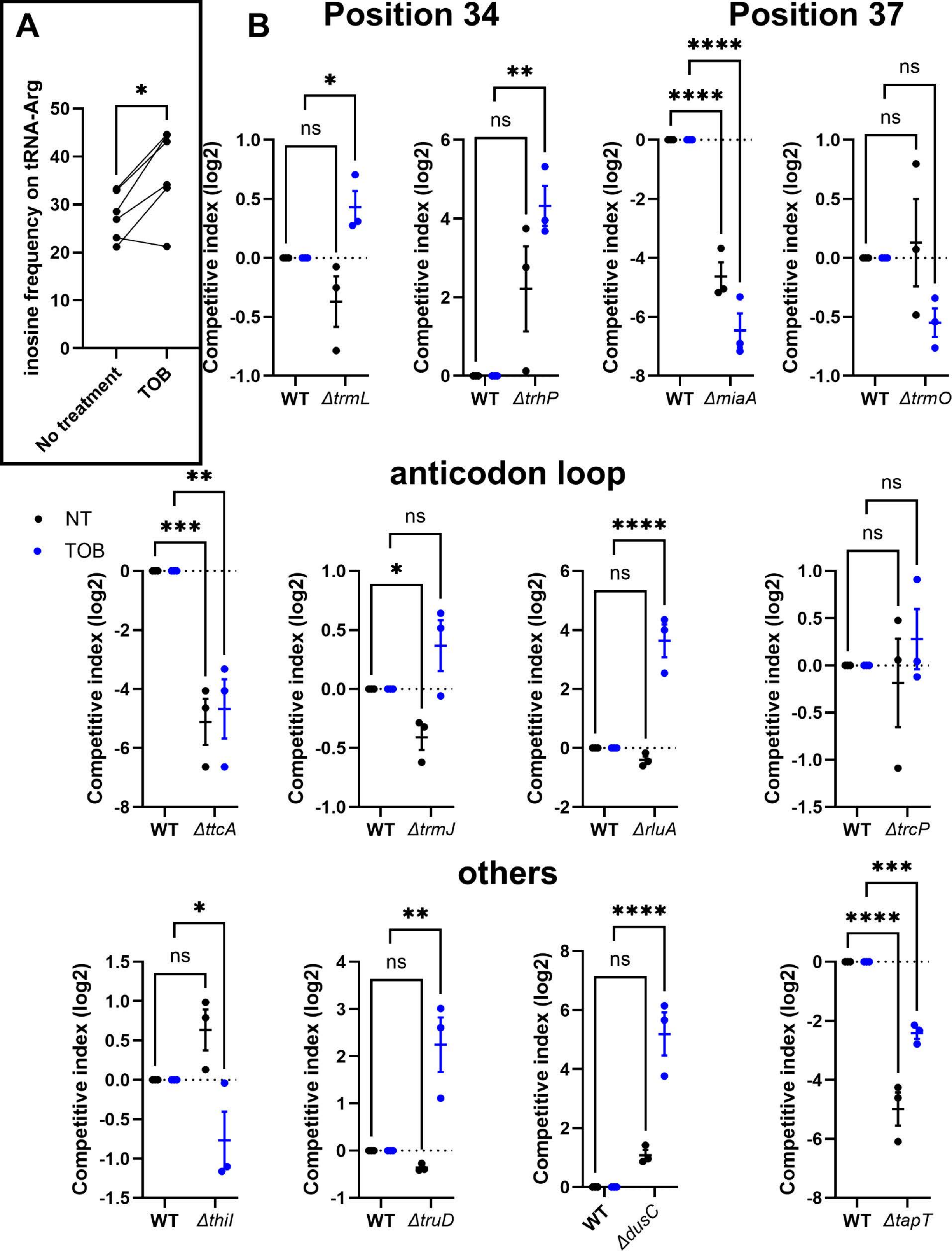

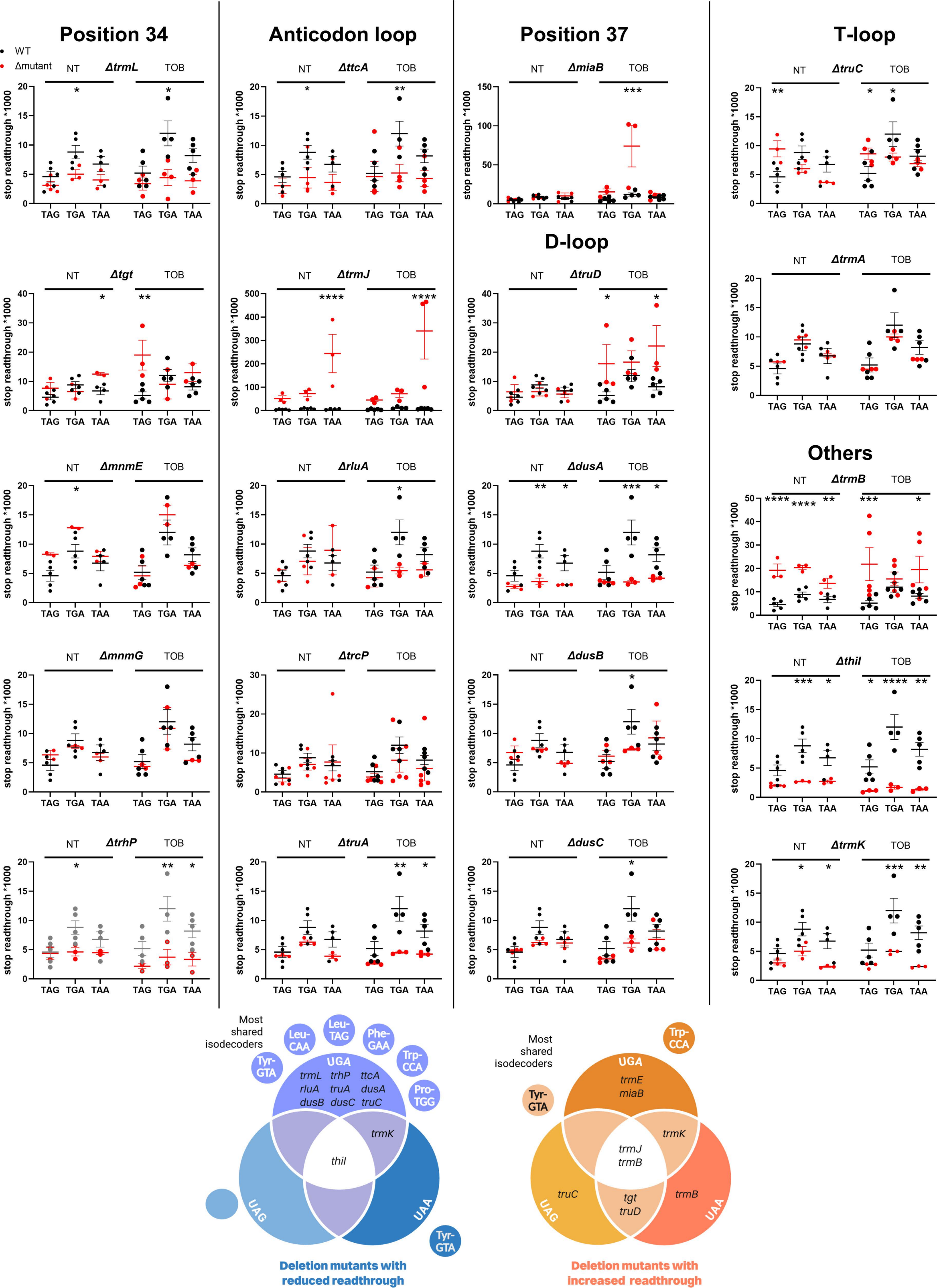

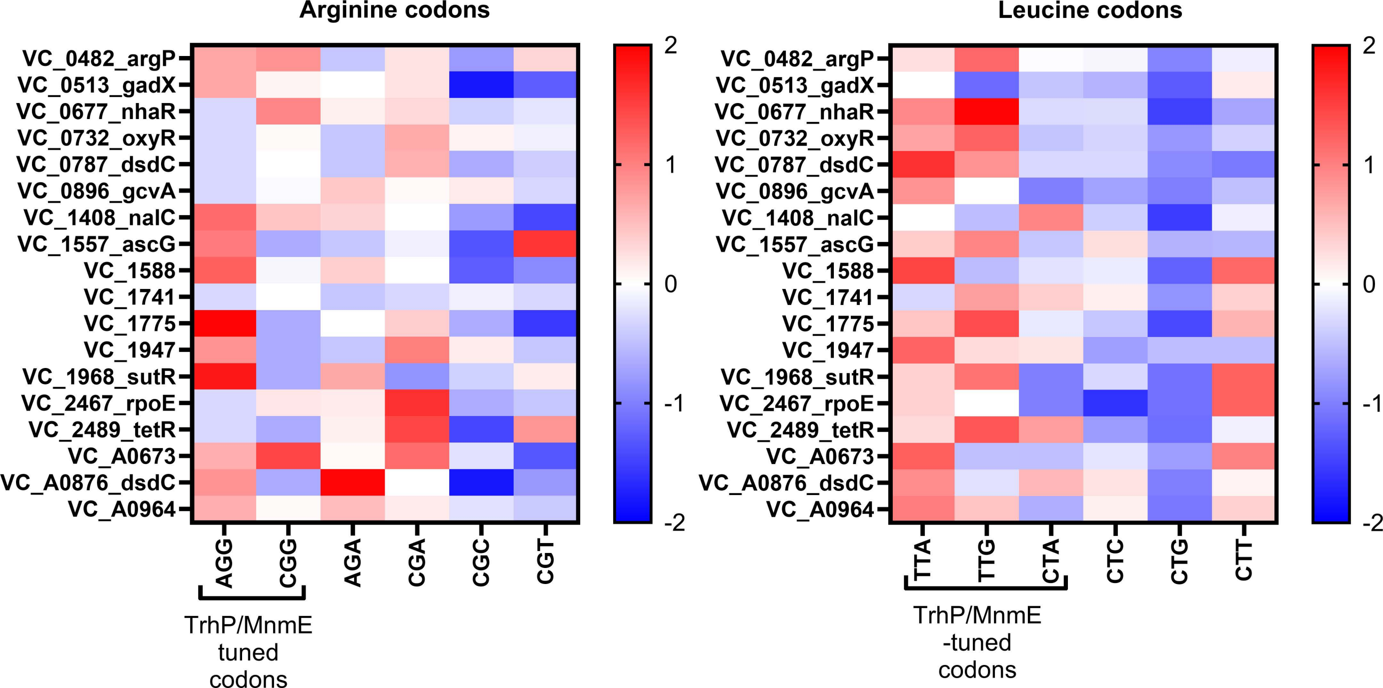

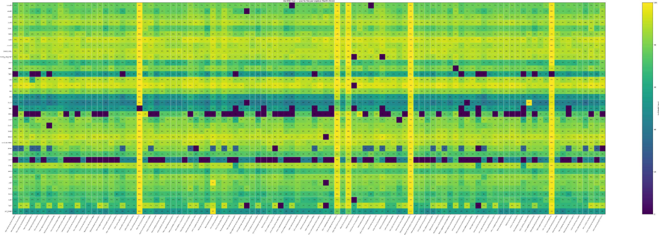
